# Role of BRCA2 DNA-binding and C-terminal domain on its mobility and conformation in DNA repair

**DOI:** 10.1101/2021.03.02.433541

**Authors:** Maarten W. Paul, Arshdeep Sidhu, Yongxin Liang, Sarah E. van Rossum-Fikkert, Hanny Odijk, Alex N. Zelensky, Roland Kanaar, Claire Wyman

**Affiliations:** Department of Molecular Genetics, Oncode Institute, Erasmus MC Cancer Institute, Erasmus University Medical Center, 3000 CA, Rotterdam, The Netherlands; Department of Radiation Oncology, Erasmus University Medical Center, 3000 CA Rotterdam, The Netherlands

## Abstract

BRCA2 is an essential protein in genome maintenance, homologous recombination and replication fork protection. Its function includes multiple interaction partners and requires timely localization to relevant sites in the nucleus. We investigated the importance of the highly conserved DNA binding domain (DBD) and C-terminal domain (CTD) of BRCA2. We generated BRCA2 variants missing one or both domains in mouse ES cells and defined their contribution in HR function and dynamic localization in the nucleus, by single particle tracking of BRCA2 mobility. Changes in molecular architecture of BRCA2 induced by binding partners of purified BRCA2 was determined by scanning force microscopy. BRCA2 mobility and DNA damage-induced increase in the immobile fraction was largely unaffected by C- terminal deletions. The purified proteins missing CTD and/or DBD were defective in architectural changes correlating with reduced homologous recombination function in cells. These results emphasize BRCA2 activity at sites of damage beyond promoting RAD51 delivery.

## Introduction

Breast cancer-associated protein 2 (BRCA2) is a required component in multi-step genome maintenance processes that are coordinated in time and place. BRCA2 knock-out is lethal in mammalian cells and defective BRCA2 causes increased sensitivity to genotoxic agents, defective DNA repair and reduced homologous recombination activity (Prakash et al., 2015; Sharan et al., 1997; Yu et al., 2000). One role of BRCA2 common to DNA break repair, DNA crosslink repair and replication fork protection, is delivery of RAD51 to sites where it is needed (Holloman, 2011; Sharan et al., 1997; Yuan et al., 1999). RAD51 is also an essential protein whose biochemical function is to form filaments on ssDNA capable of performing strand exchange reactions with homologous partners or otherwise protecting the bound DNA (Baumann & West, 1998; Heyer et al., 2010). We consider essential BRCA2 activity to involve at least (1) spatial relocation in the nucleus resulting in accumulation to sites where RAD51 is needed and (2) molecular rearrangement to release or deposit RAD51 on DNA in an active form.

Accumulation of the required proteins at the sites of DNA damage is typically defined as the appearance of foci, high local concentration of proteins, in immunofluorescence experiments. During homologous recombination the formation of RAD51 foci is considered a critical step and this has recently also introduced in clinical setting as a test for homologous recombination defects in tumors (Naipal et al., 2014). The accumulation of RAD51 into foci depends on functional BRCA2 (Yuan et al., 1999). Several studies have addressed the role of different interactors and domains of BRCA2 on foci formation after DNA damage induction (Shahid et al., 2014). The presence of Partner and Localizer of BRCA2 (PALB2) and its interaction with the N-terminus of BRCA2 is essential for the localization of BRCA2 and RAD51 to foci (Oliver et al., 2009; Xia et al., 2007; Xia et al., 2006), whereas loss of the interaction affects homologous recombination and genome stability in general (Hartford et al., 2016). In chicken DT40 cells both N-terminal interaction with PALB2 and C-terminal DBD have a role in focal accumulation of BRCA2, accumulation is fully eliminated if neither domain is present (Al Abo et al., 2014). Additionally, the interaction of the BRCA2 DNA binding domain with the small DSS1 protein is required for proper localization of BRCA2 to the nucleus and for BRCA2 and RAD51 focus formation (Gudmundsdottir et al., 2004; Kojic et al., 2005; Li et al., 2006).

Accumulation of BRCA2 and RAD51 in DNA damage induced foci necessarily requires a change in their diffusive behavior. Single particle tracking in living mouse embryonic stem (mES) cells revealed that BRCA2 diffuses as multimeric complexes bound to all detectable nuclear RAD51 (Reuter et al., 2014). Individual BRCA2 particles diffuse slowly and are transiently immobile, and this immobility increases in response to DNA damage induction (Reuter et al., 2014). Although they diffuse together, BRCA2 and RAD51 are separated at sites where they accumulate, as determined by super-resolution microscopy (Sanchez et al., 2017; Whelan et al., 2018). This suggests structural rearrangements of the complex to release RAD51. Purified BRCA2 protein shows remarkable rearrangement by RAD51, ssDNA and ssDNA (Le et al., 2020; Sanchez et al., 2017; Sidhu et al., 2020). This apparent structural plasticity is a hallmark of proteins with intrinsically disordered regions (Dunker et al., 2005; Gunasekaran et al., 2003; van der Lee et al., 2014), which we hypothesize could be relevant for BRCA2 function in cells.

BRCA2 has many predicted disordered regions, which complicate understanding the functional organization and possible dynamic rearrangement of the reported structured domains. Several crystal structures of small fragments of BRCA2 are available: C-terminal BRCA2- DSS1 complex (Yang et al., 2002)(PDB ID: 1IYJ), N-terminal BRCA2-PALB2 (Oliver et al., 2009)(PDB ID: 3EU7), Brc4 BRCA2-RAD51 (Pellegrini et al., 2002)(PDB ID: 1N0W), BRCA2 phosphopeptide-Plk1 (Ehlen et al., 2020)(PDB ID: 6GY2), Brc8-2 BRCA2-RadA (Lindenburg et al., 2020)(PDB ID: 6HQU) and a BRCA2 peptide residing in exon 12 with the C-terminal ARM domain of HSF2BP (Ghouil et al., 2020). Of these the largest crystalized segment of BRCA2 is the DNA binding domain (736 amino acids), which encompasses the helical domain, tower domain and the 3 OB folds. There are two distinct regions of BRCA2 that bind RAD51 with different consequences (Esashi et al., 2005; Galkin et al., 2005). The 8 centrally located BRC repeats bind multiple RAD51 molecules (Jensen et al., 2010). The BRC repeat- RAD51 interactions are important for localizing RAD51 into nuclear foci (Chen et al., 1999) and promoting RAD51 filament nucleation in vitro (Shahid et al., 2014). There is another, single phosphorylation-dependent, RAD51 interaction domain at the C-terminus of BRCA2 (Esashi et al., 2005). This region, the C-terminal domain (CTD) of BRCA2, which is equivalent to exon 27, has a specific role protecting replication forks (Feng & Jasin, 2017; Lomonosov et al., 2003; Schlacher et al., 2011). The C-terminal RAD51 binding site is also suggested to stimulate RAD51-mediated recombination and stabilize RAD51 filaments (Esashi et al., 2005) while the DBD in combination with DSS1 is suggested to exchange RPA for RAD51 on ssDNA (Yang et al., 2002; Zhao et al., 2015). However, the DBD is not essential for cells to survive (Edwards et al., 2008) and although it is the most conserved parts of BRCA2 (Yang et al., 2002) its exact cellular function remains elusive.

It is essential for understanding how BRCA2 works to know how the regions of BRCA2, interacting with multiple partners, contribute to its function. Here we consider the influence of the DNA-binding (DBD) and C-terminal domain (CTD) on dynamic activities of BRCA2; DNA damage induced changes in diffusion and partner-binding induced changes in protein conformation. To understand which parts of BRCA2 are responsible for dynamic localization and structural transitions we correlated the cellular phenotypes, diffusion dynamics and in vitro structural transitions for BRCA2 variants lacking either the DBD, the CTD or both. We discover that the separate domains do not play a significant role in BRCA2’s nuclear localization and diffusion dynamics or RAD51 accumulation but strongly affect protein conformational response to binding partners. The BRCA2 conformational changes correlate with cellular homologous recombination activity. We discuss the possible importance of BRCA2 for steps in the homologous recombination beyond its identified role in RAD51 delivery.

## Results

To determine the role of DBD and CTD in DNA damage repair, BRCA2 mobility and structural plasticity we created murine cell lines and purified human BRCA2 protein lacking these domains (**Figure 1A**). Mouse ES cell (mES) lines producing tagged variants of BRCA2 (full-length; ΔDBD, containing an internal deletion of amino acids 2401-3143; ΔCTD, truncated at 3143; and ΔDBDΔCTD truncated at 2401) were engineered by homozygous modification of the endogenous *Brca2* alleles and addition of a HaloTag at the end of the coding sequence (**Figure 1B, Figure 1 - supplement 1**). This allowed us to visualize BRCA2 in live or fixed cells (Los et al., 2008) and study the effect of deletions under native expression in the absence of wild type BRCA2 (**Figure 2A, Figure 3**). To analyze the role of DBD and CTD *in vitro,* we purified variants of human BRCA2 protein containing the same deletions (**Figure 1A, Figure 4 and 5)**.

**Figure 1.**
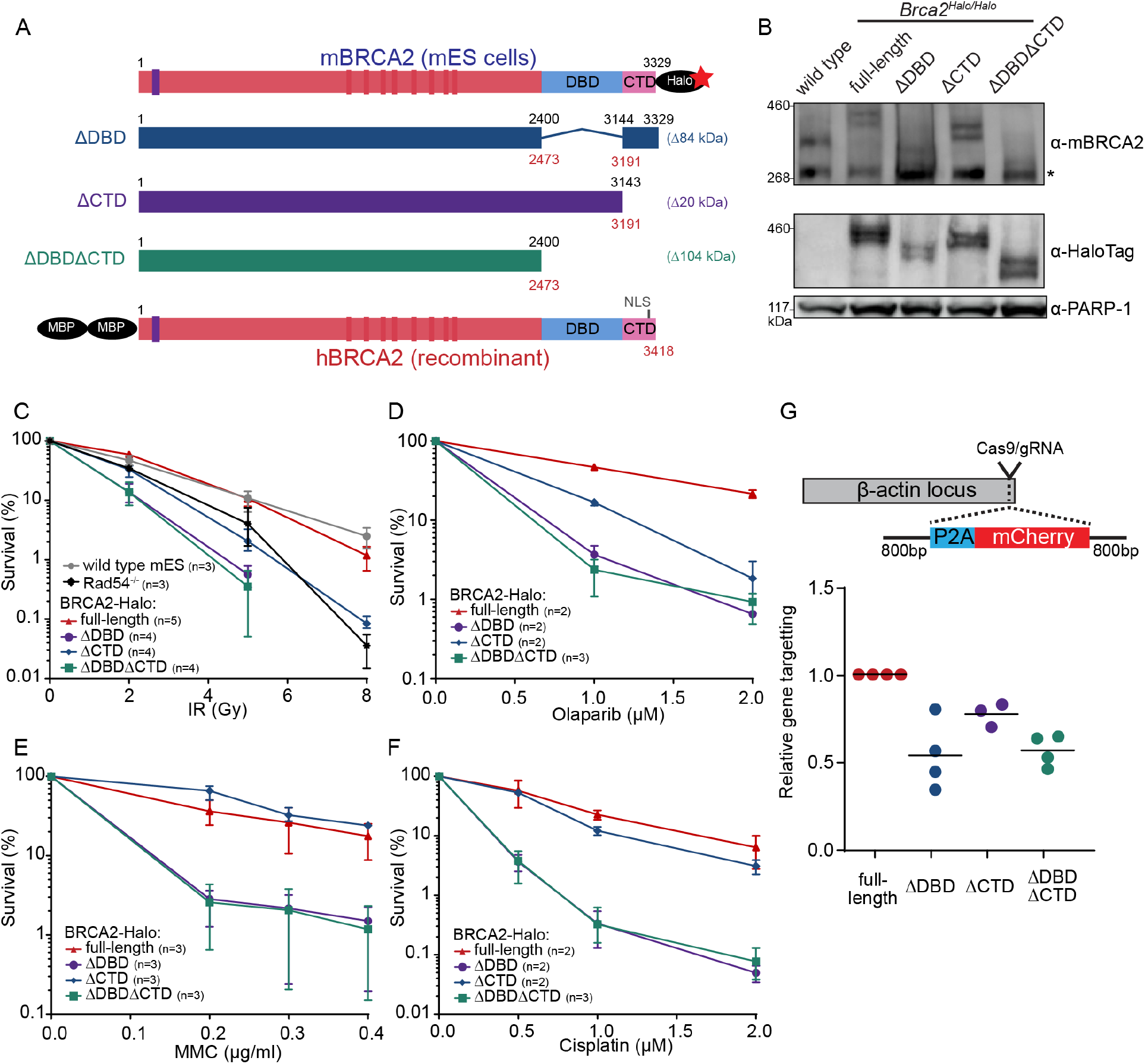
Functional analysis BRCA2 deletion variants in mouse embryonic stem cells, tagged at the endogenous locus with HaloTag. (**A**) Schematic overview of full-length mouse (top) and human (bottom) BRCA2 proteins, with key domains (DBD, CTD, NLS, BRC1-8: red bars, PALB2-binding: blue bar) and tags indicated. Deletion variants are shown in the middle. Amino acid numbers are shown in black (mouse) and red (human). Expected molecular weight decreases for the deletion variants are shown on the right. (**B**) Immunoblot of total protein extract from mES cells probed with indicated antibodies. Asterisk shows a- specific band. Validation of the cell lines by genotyping can be found in **Figure 1 – supplement 1**. (**C-F**) Clonogenetic survivals after IR, Olaparib, MMC and cisplatin treatment with the indicated doses. At 8 Gy of IR the percentage of surviving colonies of the ΔDBD- and ΔDBDΔCTD-Halo was too low to accurately determine the survivial. Error bars indicate the range of data points. (**G**) CRISPR/Cas9 based homologous recombination assay to assess the homologous recombination proficiency of the different BRCA2 mutants. mES cells are transfected with a plasmid encoding Cas9 and the specific gRNA and an repair template with the self cleaving peptide P2A and the mCherry sequence in between two homology arms. Upon proper integration of the donor sequence at the ß-Actin locus the cells will express mCherry. 96 hours after transfection cells are sorted and the frequency of mCherry positive cells is measured (**Figure 1 – supplement 2)**. To correct for difference in transfection efficiency a plasmid expressing BFP is co-transfected. The frequency of positive cells in every experimental replicate is normalized against wildype BRCA2-Halo cells.

**Figure 2.**
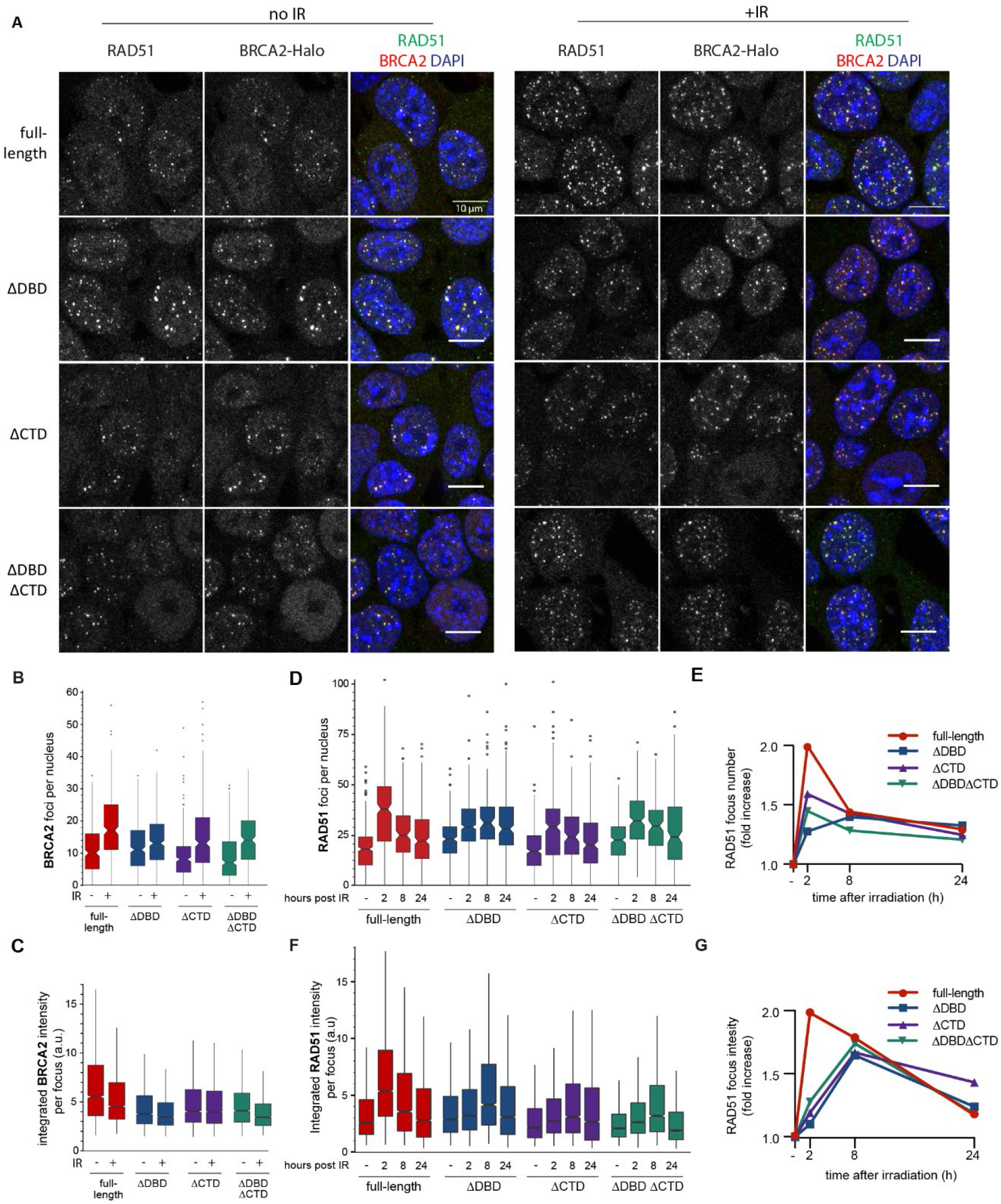
BRCA2 and RAD51 foci quantification. (**A**) Repersentative confocal images (maximum intensity projections) of BRCA2 (red) and RAD51 (green) foci in mES cells fixed 2 h after mock or 2 Gy irradiation, without pre-extraction. Scale bar 10 μm. (**B**) Quantification of the number of BRCA2-Halo (JF646) foci per nucleus of cells irradiated with 2Gy IR in cells in EdU+ without preextraction, two technical replicates, at least 250 cells per condition. (**C**) integrated BRCA2 focus intensity. (**D**) Quantification of the number of of RAD51 foci in EdU-positive cells irradiated with 2 Gy and fixed after indicated number of hours with preextraction for RAD51 immunostaining. Example images and percentage EdU+ cells per condition in **Figure 2 – Supplement 1**. Three technical replicates, at least 100 cells per condition. (**E**) Fold change of foci number with respect to untreated cells. (**F**) Integrated RAD51 intensity per focus (**G**) fold change in integrated intensity of RAD51 foci relative to untreated cells. Representative images are shown in **Figure 1 – Supplement 1**. In boxplots in C and F distribution outliers not shown.

**Figure 3:**
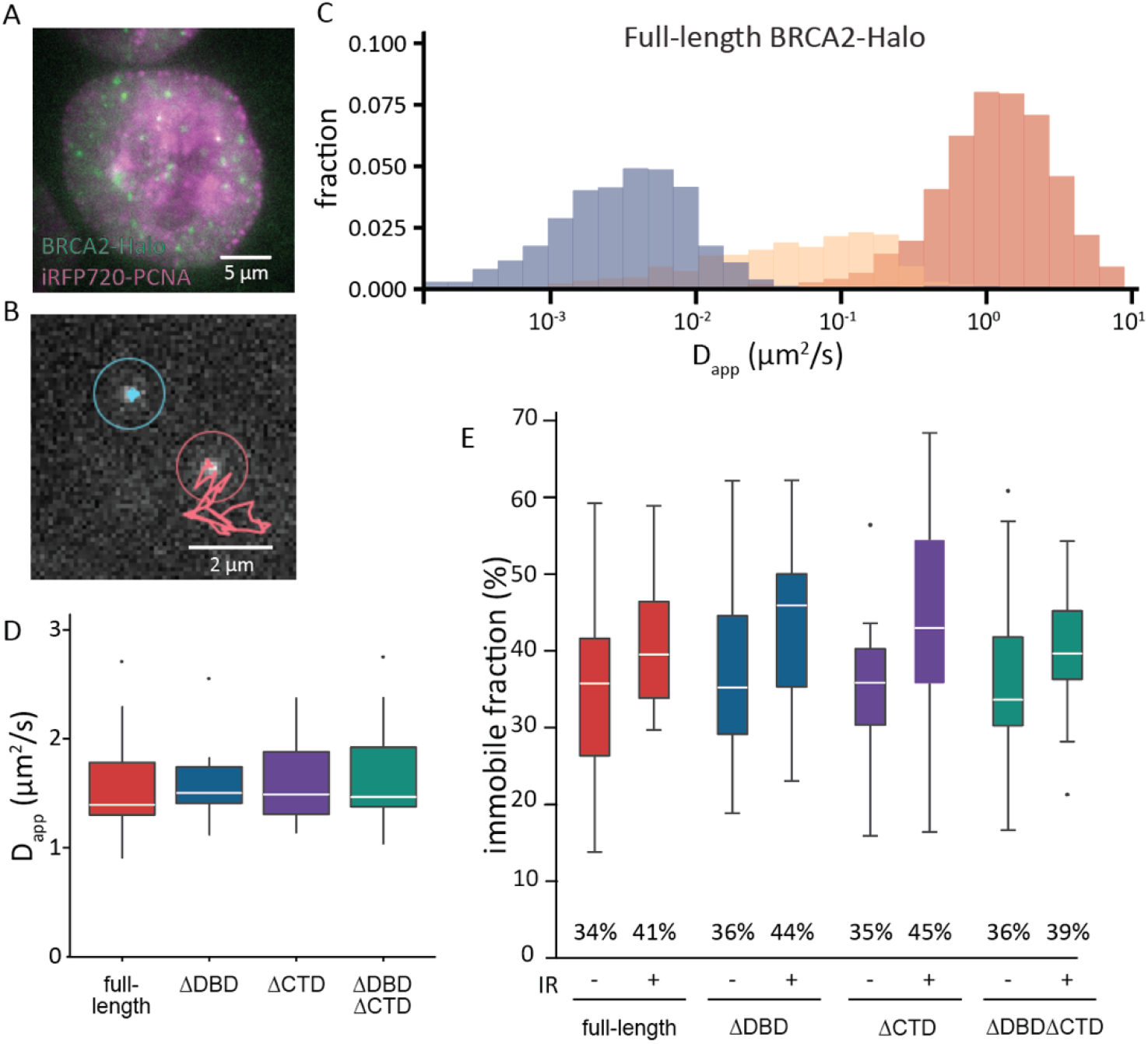
Single-particle tracking of BRCA2-HaloTag reveals immobilization of BRCA2 lacking either DBD or CTD upon DNA damage. (**A**) Wide-field image of S-phase cell visualized with iRFP720-PCNA and BRCA2-HaloTag::JF549. (**B**) Example of two tracks of BRCA2-Halo showing different diffusive behavior. (**C**) Distribution of apparent diffusion coefficients of segmented tracks (tracklets) for immobile (blue), slow (yellow) and fast (red) molecules for full-length BRCA2 in untreated cells, plots for IR treated cells and other BRCA2 variants are shown in **Figure 3 – supplement 1**. (**D**) Apparent diffusion rate of fast diffusing BRCA2 tracklets for full-length BRCA2 and indicated deletion variants. p-values (two-sided t-test) comparing full-length with deletion variants (ΔDBD, ΔCTD, ΔDBDΔCTD) respectively: p=0.953, p=0.797, p=0.593. (**E**) Immobile fraction estimated by segmentation of tracks by their immobile, slow or fast mobility (tracklets). Fraction is defined as the percentage of tracklets per cell that are immobile. Cells were imaged between 2 and 4 hours after IR treatment. p-values (two-sided t-test) comparing -/+ IR for different variants (full-length, ΔDBD, ΔCTD, ΔDBDΔCTD) respectively: p=0.08, p=0.057, p=0.02, p=0.4. Merged data from two independent experiments of at least 15 cells and about 10,000 tracks per condition. Percentages below the plot indicates the mean immobile fraction of all cells.

**Figure 4.**
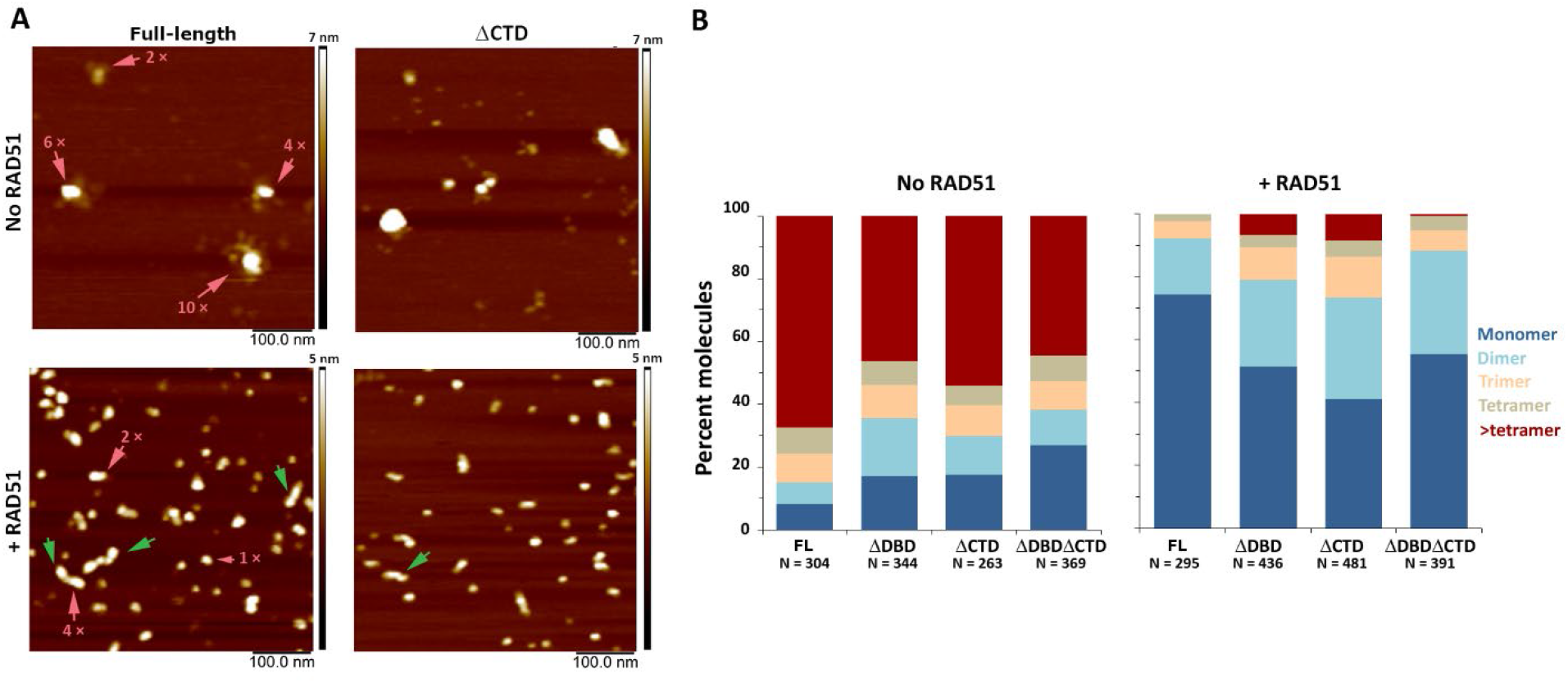
C-terminal region of human BRCA2 contributes to the formation of BRCA2-BRCA2 oligomers. (**A**) Representative SFM height images of full-length and ΔCTD BRCA2 in the presence and absence of RAD51. BRCA2 ΔCTD forms rod-like shaped assemblies, like full-length BRCA2, on interaction with RAD51. Rod-like assemblies are indicated by green arrows; pink arrows indicate multimeric assemblies, based on volume analyses. (**B**) Histograms showing oligomeric distribution of full-length BRCA2 and the C-terminal variants in the presence and absence of RAD51. The deletion of C-terminal region leads to lesser oligomeric forms than full-length BRCA2. All the experiments were performed twice with independent protein preparations, imaging and analyses. The figure is plotted from one of the duplicate data set.

### Loss of BRCA2 DBD and CTD impairs cell survival and gene targeting

To investigate whether the loss of the DBD or CTD affected sensitivity of the cells to DNA damage we performed clonogenic survival assays after treatment with different DNA damaging agents: double strand break induction by ionizing radiation (IR), replication disruption by PARP inhibitor Olaparib and DNA crosslink induction by mitomycin C (MMC) and cisplatin. Loss of CTD caused sensitization to IR (at 5 Gy up to 5-fold decrease in surviving fraction), comparable to the effect of deleting non-essential auxiliary homologous recombination protein RAD54 (Essers et al., 1997). In contrast, deletion of DBD led to a higher hypersensitization (18-fold decrease in surviving fraction or 3.6-fold more than in ΔCTD), which was not further exacerbated if CTD was also missing (**Figure 1C**). The effects of domain deletion on Olaparib, cisplatin and MMC sensitivities were similar, but notably, loss of CTD did not lead to a significant sensitization to cisplatin nor MMC (**Figure 1D-F**). Together, these results indicate that the BRCA2 DBD is important for efficient homologous recombination-mediated DNA repair while the CTD is less critical for this cellular activity.

To assay homology search and DNA strand exchange functions of homologous recombination, we performed a FACS-based gene targeting assay (Yao et al., 2017), in which Cas9 is used to induce a double strand break in the β-actin locus that is repaired by a donor plasmid including a mCherry coding sequence (**Figure 1G**). The absolute gene targeting frequency of about 5% (**Figure 1 – supplement 2**) was reduced 2-fold in BRCA2 ΔDBD and ΔDBDΔCTD cells, while CTD deletion caused an intermediate effect. Thus, DNA damage sensitivity described above correlates with this gene targeting assay indicating that BRCA2 DBD is specifically important for homologous recombination activity at two-ended DNA breaks.

### DBD and CTD are not essential for BRCA2 and RAD51 focus formation

A critical initial step of homologous recombination in cells involves BRCA2-mediated RAD51 localization to nuclear sites where it is needed, typically observed as foci in cell imaging. We focused on the response of BRCA2 and RAD51 to IR-induced DNA damage because timing of this response in wild type mES cell lines is well defined, and in contrast to genotoxic chemicals, damage induction is instantaneous and synchronous. We visualized BRCA2 protein with a bright photostable fluorophore via the HaloTag using JF646 HaloTag-ligand (Grimm et al., 2015) combined with RAD51 immunofluorescence (**Figure 2A**). As the DBD of BRCA2 binds DNA in vitro (Yang et al., 2002), it might contribute to BRCA2 localization and/or retention at the sites of damage. However, we observed formation of both spontaneous and IR-induced nuclear BRCA2 and RAD51 foci in all three BRCA2 deletion variants, where BRCA2 and RAD51 foci appear to overlap to a large extent (**Figure 2A**).

### DBD and CTD affect the amount of RAD51 and BRCA2 at repair sites

Absence of a clear qualitative effect on foci formation was unexpected, so we performed further systematic quantification of fluorescence of BRCA2-Halo-JF646 and RAD51, by immunofluorescence, in fixed cells. Only the CTD deletion affected BRCA2 foci, in the absence of induced DNA damage (background) there was a reduction in their number compared to full-length BRCA2 (20% reduction) (**Figure 2B**). Upon irradiation, the number of BRCA2 foci increased in all deletion variants, and the difference in fold increase compared to cells producing full-length BRCA2 was either small or absent. The effect of DBD and CTD deletion on the intensity of background BRCA2 foci was much more pronounced (1.5 fold reduction) (**Figure 2C**). Interestingly, in all cell lines IR-induced increase in the number of foci was accompanied by a decrease in focus intensity, suggesting that BRCA2 re-localises from the background to the IR-induced foci, but this effect was suppressed in the deletion variants (only 7% reduction in ΔDBD compared to 30% in cells expressing full-length BRCA2).

We further analyzed RAD51 focus formation and resolution over 24 hours after IR treatment (**Figure 2D,E, Figure 2 – Supplement 1**). As with BRCA2 foci, the number of RAD51 foci increased in both deletion variants and the control cells. Consistent with our previous observations in wild type ES cells, the number of RAD51 foci peaked 2 hours after IR then greadually decreased over time reaching near-background levels at 24 hours (**Figure 2D,E**). For the ΔDBD cells the number of foci increased but did not decrease over time, remaining high at 24 hours. The return to background number of foci was suppressed to a lesser extent in ΔCTD and the double mutant. The effect of DBD and CTD deletion on RAD51 focus intensity dynamics was also pronounced. In the control cells, changes in RAD51 foci intensity paralleled changes in their number: peaking at 2 hours, decreasing gradually thereafter (**Figure 2F**). In all three deletion variants, focus intensity increase was absent (ΔDBD); reduced (1.2-fold increase compared to 2-fold in control); or delayed (peak at 8 hours compared to 2 hours in control) (**Figure 2F,G**). Taken together, these results show that the deletion mutants of BRCA2 do accumulate RAD51 proteins to IR-induced lesions, however, less RAD51 accumultes and its turnover is supressed.

### DBD and CTD are not essential for BRCA2 mobility response to DNA damage

Previously we used single particle tracking (SPT) to determine diffusive behavior of BRCA2-GFP and observed a mobile fraction, which diffused slower than expected due to frequent transient interactions, and an immobile fraction, which increased upon induction of DNA damage (Reuter et al., 2014). Interaction between the DBD and DNA could be responsible for both restricting the diffusion of the mobile BRCA2 complexes and for immobilization upon DNA damage. To test this hypothesis, we performed SPT analysis of BRCA2 deletion variants labelled with JF549 via the HaloTag. The increased photostability and brightness of the JF549 fluorophore compared to GFP allowed us to follow the mobility for extended periods of time and at increased frame rate (2000 vs 200 frames, at 33 vs 20 fps, for BRCA2-HaloTag-JF549 and -GFP, respectively). To identify cells in S-phase we used PCNA fused with iRFP720 (**Figure 3A**). We tracked several hundred BRCA2 particles per nucleus where individual tracks appear as mobile or immobile (**Figure 3B**) and sometimes switch behavior. The diffusive behavior of BRCA2 was quantified using a recently developed deep learning algorithm to segment tracks into parts (tracklets) with different mobile states (Arts et al., 2019). An apparent diffusion constant was extracted for each class of segmented tracklets. This revealed different populations of BRCA2 molecules (**Figure 3C**), one with a low apparent diffusion coefficient between 0.001 and 0.01 μm^2^/s, which we consider immobile, a second fraction of slow mobile molecules with an apparent diffusion coefficient between 0.01 and 0.1 μm^2^/s and a third fraction mobile molecules with an average diffusion rate of 1.5 μm^2^/s — consistent with our previous results tracking BRCA2-GFP in mES cells (Reuter et al., 2014). Also comparable to our previous work, the fraction of immobile molecules, ~34% in untreated cells, (**Figure 3C**) increased to 41 % after DNA damage induction by IR (**Figure 3E**).

The BRCA2 deletion variants all had a similar apparent diffusion coefficient, mobile molecules diffuse with a rate like the full-length protein (**Figure 3D, Figure 3 – supplement 1**). The increase in immobile tracklets after IR for the variants BRCA2 ΔDBD and ΔCTD was similar to full-length, indicating that these domains separately are not essential for this change in mobility (**Figure 3E**). However, the immobile fraction for BRCA2 ΔDBDΔCTD, missing both regions, did not increase after IR as much as the others (3% increase compared to 7-10%, **Figure 3D**). Thus, either the DBD or CTD is sufficient for BRCA2 mobility changes in response to IR but protein missing both of these domains reduces this response. As deletion of single domains, ΔDBD or ΔCTD, did cause increased sensitivity to DNA damaging agents (**Figure 1**) we can conclude that diffusion changes in response to DNA damage were not sufficient to assure cell survival or proper homologous recombination activity.

### Architectural rearrangement of BRCA2 variants

Our observations so far indicated that BRCA2 function needed for DNA damage survival and homologous recombination includes activities beyond immobility. We considered that homologous recombination DNA damage response requires dynamic interaction between BRCA2 and RAD51 at a scale not evident in our (live) cell imaging. Although BRCA2 and RAD51 diffuse together in the nucleus, they are separated at the sites of DNA damage requiring a local change in RAD51 and BRCA2 interaction (Reuter et al., 2014; Sanchez et al., 2017). Previously, we have defined distinct architectural changes in full-length BRCA2 upon association with RAD51 and single-stranded DNA (ssDNA) as evidence for such a dynamic interaction (Sanchez et al., 2017; Sidhu et al., 2020). To correlate BRCA2 architectural changes with in vivo functions, we purified variants of human BRCA2 and deletion variants analogous to those tested in mES cells (**Figure 1A**). Scanning force microscopy (SFM) imaging revealed that purified BRCA2 exists as a mixture of particles of varying size (multimeric form) and shape (compact to extended). As described in the section below, these features were quantified for all individual proteins/complexes from SFM images by measuring volume and solidity (Sanchez et al., 2017; Sidhu et al., 2020). The architectural plasticity of full-length BRCA2 is evident in the change in distribution of these features upon addition of binding partners, RAD51 or ssDNA oligonucleotides (Sanchez et al., 2017; Sidhu et al., 2020).

### CTD and DBD contribute to BRCA2 self-oligomerization

All three BRCA2 deletion variants exist as a distribution of irregular oligomeric molecules as previously observed for the full-length protein (**Figures 4A, B and Figure 4 – supplement 1**) (Sanchez et al., 2017; Sidhu et al., 2020). In the conditions used here the majority (70%) of full-length BRCA2 was present as assemblies larger than tetramers (**Figure 4B: left panel**). The C-terminal deletion variants show reduced oligomerization for all the variants with 46% BRCA2 ΔDBD, 54% BRCA2 ΔCTD and 44% BRCA2 ΔDBDΔCTD present as assemblies larger than tetramers (**Figure 4B: left panel and Figure 4 – supplement 1**).

Decrease in large oligomers coincides with an increase in the monomer population from <10% in fulllength to ~30% in BRCA2 ΔDBDΔCTD (**Figure 4B: left panel**), indicating that both DBD and CTD contribute to BRCA2 interactions with itself, at least in the absence of other binding partners.

### DBD and CTD are needed for ssDNA and RAD51 induced architectural rearrangement of BRCA2

Both RAD51 and ssDNA induce notable changes in BRCA2 architecture. In the presence of RAD51, BRCA2 forms compact structures. In contrast ssDNA induces the opposite, characteristic extended forms of BRCA2 become prevalent (Sanchez et al., 2017; Sidhu et al., 2020) (**Figure S1**). Full-length BRCA2 is present in oligomeric complexes with extensions, in a bimodal distribution of solidity - a measure for the compactness of the protein complexes-with peaks at 0.75 and 0.95 (**Figure 5A and B**). Incubation with ssDNA shifts the solidity distribution to a single peak at 0.7 (**Figure 5: FL BRCA2 + ssDNA**). The BRCA2 C-terminal deletion variants had fewer extended molecules to begin with, as solidity shows a peak distribution around 0.9 for all the 3 variants; ΔDBD, ΔCTD and ΔDBDΔCTD (**Figure 5 and Table S3**). In striking contrast to the full-length BRCA2, interaction with ssDNA did not change the distribution of oligomers and shape (solidity) for any of the deletion variants (**Figure 5, Figure 5 – supplement 1**). Both the DBD and CTD have to be present for BRCA2 to undergo the conformational change associated with ssDNA interactions.

**Figure 5.**
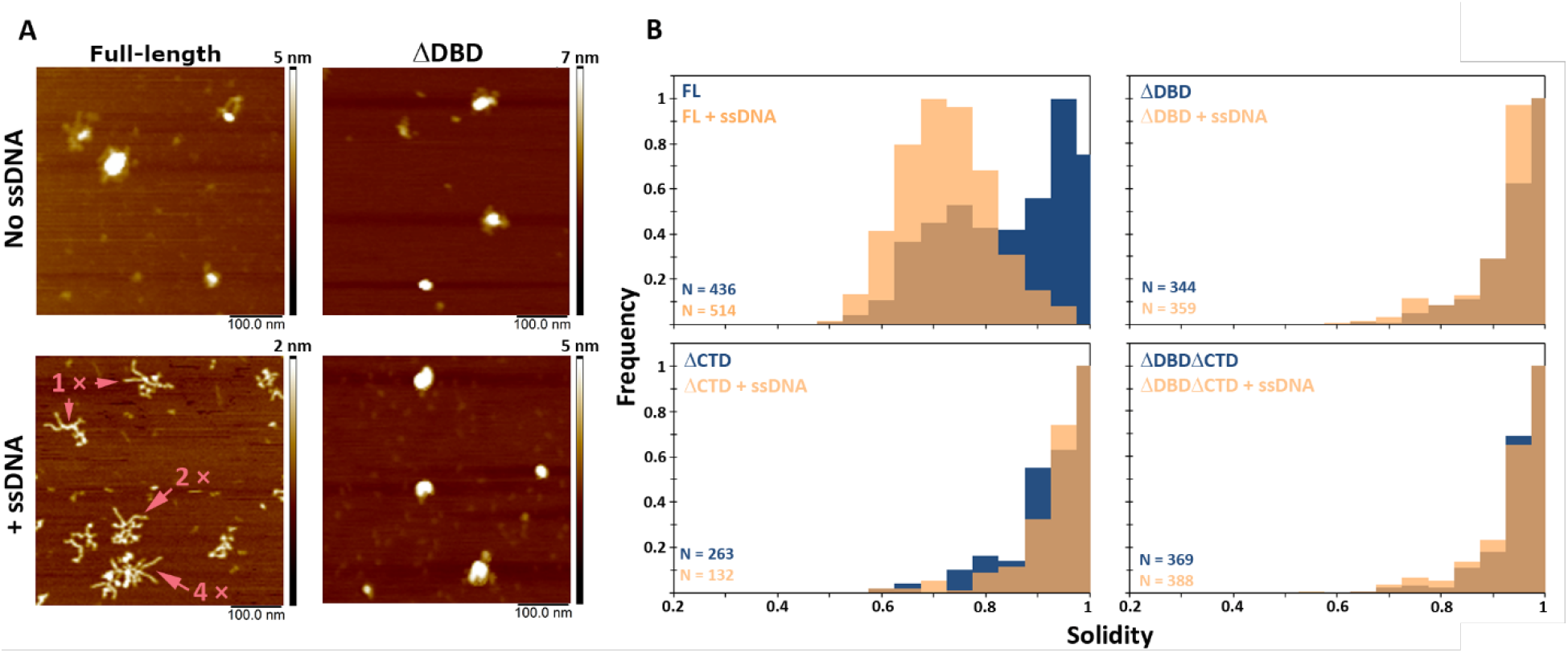
C-terminal region of BRCA2 is essential for conformational rearrangement on interaction with ssDNA. (A) Representative SFM height images of full-length BRCA2 and BRCA2 ΔDBD in the presence and absence of ssDNA. Full-length BRCA2 rearranges into extended molecular assemblies on interaction with ssDNA, however BRCA2 ΔDBD and other C-terminal constructs do not show conformational change. Pink arrows indicate the oligomeric volume of the particle with respect to BRCA2 monomer (B) Distribution of full-length BRCA2 and the C-terminal deletion constructs with respect to their oligomerization and solidity. Full-length-BRCA2 rearranges to form extended dimers and tetramers on interaction with ssDNA, whereas the deletion constructs do not show change in their distribution. All the experiments were performed twice with independent protein preparations, imaging and analyses.

Upon incubation with RAD51, full-length BRCA2 assemblies become largely monomeric (74%) and adopt a more regular compact conformation, with 33% having a rod-like shape (major to minor axis ratio >1.5) (**Figures 4A, B, Figure 4 – Supplement 1,2 and Table S4**). Purified BRCA2 binds 6 RAD51 monomers in conditions similar to ours (Jensen et al., 2010; Liu et al., 2010). Our volume-based monomer designation refers to one BRCA2 plus RAD51, as the theoretical volume of one BRCA2 and 1-6 RAD51 molecules falls in the range BRCA2 monomer (see materials and methods for detailed description of volume analysis). All deletion variants also become largely monomeric upon interaction with RAD51, but to a lesser extent than the full-length BRCA2 (40-55% for variants vs 74% for full-length). However, all the variants included about one-third of the complexes as dimers: BRCA2 ΔDBD (28%), BRCA2 ΔCTD (32%) and BRCA2 ΔDBDΔCTD (33%), which was more than the full-length BRCA2 (18%) (**Figure 4B, C: right panel**). Only BRCA2 ΔCTD-RAD51 formed rod-shaped assemblies similar to full-length BRCA2 (**Figures 4B and Table S4**). Removing either the DBD, CTD or both, reduced BRCA2 oligomerization and to a lesser extent reduced RAD51 induced changes in oligomerization and architecture (**Figure 4 supplement 1 and 2**). The effect of the BRCA2 DBD and CTD domain deletions on cellular response to DNA damage (**Figure 1**) and their effect on the architecture of BRCA2 and its complexes with RAD51 and ssDNA did correlate. The architectural changes we defined here may report on important BRCA2 cellular functions.

## Discussion

Here we investigated the role of DBD and CTD on the diffusive behavior and ligand-induced structural plasticity of BRCA2. We correlated these observations with functional consequences of deleting these domains, individually and in combination, in living cells. Our panel of isogenic precision-engineered cell lines allowed us to label endogenously expressed BRCA2 directly via the HaloTag. We found that substantial reduction in DNA damage resistance, especially in DBD-deficient cells, was accompanied by only subtle changes in dynamics and localization of BRCA2. In contrast, the ability of purified recombinant BRCA2 to undergo structural rearrangements was strongly affected by DBD or CTD deletions.

Despite their adjacent location in the C-terminal part of BRCA2 (and frequent simultaneous loss due to human cancer-predisposing mutations), DBD and CTD are functionally distinct. CTD, although much shorter that the DBD, performs several distinct functions: cell cycle-controlled phosphorylation-dependent stabilization of RAD51 filament in vitro; replication fork protection from excessive nucleolytic processing; and nuclear import. In mouse BRCA2 an additional nuclear localization signal is present at the N-terminus, but in the human protein there is no such redundancy, which exaggerates the consequence of even short C-terminal truncations, because these produce human BRCA2 that cannot localize to the nucleus (Sarkisian et al., 2001; Spain et al., 1999). Controlled deletions, including the internal DBD deletion, allowed us to avoid some of the confounding effects complicating previously used mutant or patient cell models.

The DBD is the evolutionarily defining part of BRCA2, conserved from fungi to humans, but its function is less defined than that of other, “younger” BRCA2 regions. Information on DBD function focuses on its interaction with an intrinsically disordered acidic protein DSS1. Our findings reinforce the notion that despite its deep phylogenetic roots, the DBD is not what makes BRCA2 essential for general viability of animal cells. Absence of the DBD leads to significant sensitization to DNA interstrand crosslinks, PARP inhibitor and radiation, not further exacerbated by additional CTD deletion (**Figure 1**). The role of DSS1 interaction remains puzzling: On the one hand, it is as conserved the DBD itself, was shown to be required for BRCA2 stability and intracellular localization (Li et al., 2006); mutations in DSS1 binding phenocopy BRCA2 deficiency, as does DSS1 depletion (Zhao et al., 2015). But, on the other hand, in fungi DSS1 is only required for DNA repair when DBD is present (Kojic et al., 2005). Similarly, in human cells homologous recombination could be partially restored in BRCA2-deficient cells by complementation with variants lacking the DBD (Edwards et al., 2008; Siaud et al., 2011). Our results also show that cells expressing BRCA2 ΔDBD retained ~50% of homologous recombination activity (**Figure 1G**). We also found that DBD contributes little to the characteristic constrained diffusion we described previously. Binding of the DBD to DNA could explain slow diffusion and frequent immobilization of BRCA2 — with higher frequency after damage induction. But the effect of loss of DBD on diffusive activity was small and comparable to loss of CTD. Therefore, we conclude that the DBD does not have a significant role in this context. In the BRCA2-containing complex DNA binding activity could be redundantly supplied by its interactors. For example, several BRCA2-bound RAD51 molecules provides alternative DNA interactions interfaces (Jensen et al., 2010; Reuter et al., 2014; Sanchez et al., 2017).

In contrast to the DBD, the CTD is a recent vertebrate addition to BRCA2. In our assays its deletion resulted in no phenotype (interstrand crosslink survival), intermediate phenotype (radiation and PARPi survival, homologous recombination assay, RAD51 focus number) or the same effect as DBD deletion (RAD51 focus intensity). Except for BRCA2 oligomerization (**Figure 4B**) deleting both domains did not result in an additive effect. It is possible that CTD deletion we created encroaches on or disturbs the structure of the DBD, and (some of) the functional consequences we attribute to the CTD deletion result from collateral damage to the DBD. The strongest argument against this is the clear separation of functions between the domains in the interstrand crosslink survival assays (**Figure 1E, F**) where CTD deletion has no effect. This finding is also suggesting that the described fork protection and RAD51 filament stabilization functions of the CTD are not essential for DNA crosslink repair. This is at odds with studies that describe the role of fork protection in crosslink repair, and in particular with previous findings on ES cells with different CTD-disrupting Brca2 alleles (Atanassov et al., 2005; Donoho et al., 2003; Marple et al., 2006). Details of the genomic engineering strategies could account for the differences: for example, the more widely used Brca2 lex1/lex2 cells are compound heterozygote, with a larger deletion in one of the alleles (Morimatsu et al., 1998). Another unexpected observation was that CTD deletion blocked structural rearrangements of BRCA2 upon interaction with ssDNA as efficiently as the DBD deletion. One possibility is that N-C terminal interactions that contribute to oligomerization of BRCA2 (Le et al., 2020) also effect BRCA2 structure and thereby influence the BRCA2-ssDNA interaction.

We observed RAD51 focus formation in some cell lines that were however deficient in homologous recombination at two-ended double-strand breaks. Despite strong sensitization to radio- and chemotherapeutic agents, only careful quantification of the numbers and intensity of RAD51 foci at multiple timepoints after radiation revealed subtle differences in the deletion variants. The reduced intensity of RAD51 foci in cells lacking DBD and CTD indicates that the repair process is delayed or reduced at some point beyond delivery of RAD51 by BRCA2 to the sites of damage. This suggests that RAD51 foci quantification, although useful to identify more homologous recombination deficient samples than BRCA1/2 mutations, could occasionally return false positive results for homologous recombination function at two-ended breaks. In and of itself this observation is not surprising as there will be steps important for homologous recombination function downstream of RAD51 focus formation. However, it does raise a note of caution in the context of employing the RAD51 focus formation assay in pre-clinical and clinical settings with the aim to find BRCAness phenotypes. Additional homologous recombination markers may need to be explored to overcome this limitation.

The DBD and CTD of BRCA2 did markedly affect protein architecture and conformational changes in response to binding partners. These domains contribute to oligomerization, when we remove them, in DBD and CTD deletion variants, the BRCA2 population was less oligomeric. This is an agreement with a recent study where interaction of N and C-terminal fragments of BRCA2 is indicated to contribute to oligomerization of BRCA2 (Le et al., 2020). However, in our study the oligomeric forms induced by RAD51 binding remain unchanged, which is likely mediated by interaction with the intact BRC repeats. The characteristic conformational change of irregular compact particles-to-extended architecture of fulllength BRCA2 in response to ssDNA was severely impaired in all the investigated deletion variants. Together, the inability of the deletion variants to rearrange in vitro in the presence of ssDNA coupled with the impaired homologous recombination in vivo suggests DBD and CTD interactions of BRCA2 are important for optimal BRCA2 activity at the sites of damage. Similar regulatory function is reported for other proteins in that interact with BRCA2 such as DSS1, which also affects the conformation of BRCA2 (Le et al., 2020). Suggesting that regulation of RAD51 by BRCA2, is affected by conformational rearrangement of BRCA2 and is mediated at different levels by self-interaction of BRCA2 and its interaction partners.

Comparing all the molecular endpoints we analyzed (diffusion, foci, architecture) we conclude that although none correlated perfectly with the functional outcomes (survival and recombination assay), the magnitude of the effect on architectural plasticity was the closest reflection. We are only starting to tease apart the relationship between structural plasticity and cellular function. BRCA2 function may depend not so much on the existence of one structural form or another but on the lifetime of specific conformations affected by its interactors and local chromatin organization, parameters that will need to be quantified.

## Materials and methods

### Plasmids for cell experiments

Plasmids containing gRNAs and spCas9 were derived from px459 (Ran et al., 2013). As described in Zelensky et al. (2017), selected gRNA sequences were incorporated into the px459 vector (Ran et al., 2013) by digestion of the vector with AflIII and XbaI. The resulting two fragments (vector backbone and restriction fragment) were separately purified from gel. The resulting restriction fragment was used as template for two PCRs with overhanging primers containing the required gRNA sequence (see Table S1). Using Gibson assembly, the two fragments and the digested vector backbone were assembled and transformed in E. coli (DH5 alpha). The correct integration of the gRNA sequence in the isolated plasmid was validated by Sanger sequencing. For incorporation of two gRNA in a single plasmid, px459 was modified to contain two U6 promotors and gRNA sequences separated by a short spacer.

The donor template for C-terminal tagging of BRCA2 with HaloTag was derived from the plasmid that was used to make BRCA2-GFP knock-in cell lines (Reuter et al., 2014). This plasmid contains 3’ and 5’ homology arms (6.6 and 5.4 kb homology) for integration of the construct at the BRCA2 locus. The GFP sequence was removed by restriction digest and replaced with the HaloTag sequence by Gibson Assembly. The HaloTag sequence was obtained by PCR from pENTR4-HaloTag (gift from Eric Campeau, Addgene #29644). The donor plasmids for the ΔCTD, ΔDBDΔCTD-HaloTag contain a 6kb homology arm upstream of the deletion, while the downstream homology arm was identical to the full-length construct (See **Figure 1 – supplement 1**). The ΔDBD donor construct was made by introducing the coding sequence from exon 27 of mouse BRCA2 excluding the stop codon in the ΔDBDΔCTD-HaloTag donor construct.

The PiggyBac iRFP720-PCNA construct was generated using Gibson assembly, by inserting the iRFP720 sequence (Shcherbakova & Verkhusha, 2013) and hPCNA sequence (Essers et al., 2005), which includes an additional NLS sequence in a PiggyBac vector (Zelensky et al., 2017) containing a CAG promotor and PGK-puro selection cassette. iRFP720 was obtained by PCR from iRFP720-N1 (gift from Vladislav Verkhusha, Addgene #45461).

### Cell culture

Wildtype (IB10, subclone of E14 129/Ola (Hooper et al., 1987)) and *Rad54-/-* mouse ES cells (Essers et al., 1997) were cultured on gelatinized plates (0.1% porcine gelatin (Sigma)). The culture media consisted of 50% DMEM (High-Glucose, Ultraglutamine, Lonza), 40% BRL conditioned medium, 10 % FCS supplemented with non-essential amino acids, 0.1 mM β-mercaptoethanol, pen/strap and 1,000 U/ml leukemia inhibitory factor (mouse).

For imaging, cells are seeded in 8-well glass-bottom dishes (Ibidi) which were coated with 25 μg/ml laminin (Roche) for at least one hour. About 30 000 cells in 300 μl medium were plated per well the day before the experiment. Cells treated with IR were irradiated in an Xstrahl RS320 X-Ray generator (195.0 kV and 10.0 mA) at the indicated dose.

### Generation of HaloTag knock-in cell lines

15 μg of circular donor plasmid and 15 μg of px459 containing Cas9 and the indicated gRNA(‘s) were electroporated into 10^7^ IB10 mouse ES cells. About 24 hours after electroporation cells were put on selective medium containing 200 μg/ml G418 (Formedium). Medium was refreshed regularly, at least once every second day, and after 8-10 days colonies were picked into a gelatin coated 96 well plate. After 2 days cells in the 96-well plate were split and part of the cells were incubated in lysis buffer (50 mM KCl, 10 mM Tris-HCl pH 9, 0.1% Triton X-100, 0.15 μg/ml proteinase K) at 50°C for 1 hour. After inactivation of proteinase K at 95°C for 10 minutes, cell lysates were diluted and 5 μl was used for genotyping PCR using MyTaq DNA polymerase (Bioline) with indicated DNA primers. Selected (homozygous) clones were expanded and knock-ins were validated by western blot as described in Reuter et al. (2014)on a 5% SDS-PAGE gel using rabbit polyclonal anti-BRCA2 (Abcam, ab27976). Anti-HaloTag (mouse monoclonal, Promega G9211) blot (**Figure 1B**) was run on a NuPAGE 3-8% Tris-acetate gel (Invitrogen). From selected clones genomic DNA was isolated using phenol extraction and genotyping PCRs were done as described above.

### Clonogenic survivals

For clonogenic survivals, between 100 and 15 000 cells were seeded in gelatin-coated 6-well plates. The next day about 16 hours later, cells were treated at the indicated doses with ionizing radiation (IR) or the next day incubated for 2 hours with mitomycin C (Sigma-Aldrich, M503) or for 24 hours with olaparib or cisplatin after which the cell media was refreshed. 5-7 days after treatment the cells were stained with Coomassie Brilliant Blue and manually counted.

### Homologous recombination assay

The AAV_Actb HR donor plasmid used for the Cas9-stimulated HR gene targeting assay was a gift from Hui Yang (Addgene plasmid #97317) and consisted of 800bp homology arms targeting the β-actin locus with the P2A-mCherry sequence between the homology arms, as described in Yao et al. (2017). The gRNA targeted the same sequence (agtccgcctagaagcacttg) as in the original paper and was cloned into px459 (Ran et al., 2013) as described above for the other Cas9/gRNA constructs.

250 000 cells were seeded in 24 well plates and were directly transfected with Lipofectamine 3000, using manufacturer’s instructions, using 0.5 μg donor, 0.5 μg px459 and 0.1 μg pGB-TagBFP2. 24 hours after transfection the medium was refreshed. Cells were measured by flow cytometry (BD LSRFortessa) 4 days post transfection to determine the efficiency of homologous integration of the P2A-mCherry sequence. After gating for single live cells, transfected cells were gated based on BFP2 expression and subsequently the percentage of mCherry positive cells was determined (**Figure 1 – supplement 2**). We confirmed by transfection of the donor plasmid without gRNA and Cas9 that positive cells were not due to background expression of the donor plasmid.

### Immunofluorescence

Cells were grown in 8-well glass-bottom dishes (80826, Ibidi) as described above. When indicated BRCA2-HaloTag cells were incubated with 250 nM JF549 or JF646 HaloTag ligand. Cells were washed with PBS and fixed in 4% PFA in PBS for 15-20 minutes. Cells were washed with 0.1% Triton in PBS and blocked in blocking buffer (PBS with 0.5% BSA and 1.5 g/L glycine). Primary antibodies are diluted in blocking buffer and incubated with the sample for 2 hours at room temperature. Slides are washed in PBS with 0.1% Triton and subsequently incubated with secondary antibodies in blocking buffer for 1 hour at room temperature. Cells were washed in PBS and DNA was labelled by incubation with DAPI (0.4 μg/ml).

For RAD51 focus quantification on replicating cells specifically, 15 minutes before fixation cells were treated 20 μM EdU in the medium at 37 °C. Cells were washed with PBS, pre-extracted for 1 minute (300mM sucrose, 0.5% Triton X-100, 20 mM Hepes KOH (pH 7.9), 50 mM NaCl, 3 mM MgCl2), washed with PBS and directly fixed in 4% PFA in PBS. For EdU Click-chemistry labelling cells were washed with 3% BSA in PBS, permeabilized with 0.5% Triton in PBS for 20 minutes. After another wash with 3% BSA samples were incubated in home-made click-labelling buffer (50 mM Tris, 4 mM CuSO4, 10 mM Ascorbic acid and 60 μM Atto568 azide (ATTO-TEC GmbH)) for 20 minutes in the dark. For BRCA2 focus quantification in replicating cells Atto488 (ATTO-TEC GmbH) was used instead. Subsequently immunofluorescence was performed as described above and DNA was stained using DAPI.

### Confocal microscopy

Confocal images were acquired at a Zeiss Elyra PS1 system with additional confocal scan unit coupled to an Argon laser for 488nm excitation (Alexa 488) and additional 30mW 405nm (DAPI), 10mW 561nm (CF568, JF549) and 633nm (Alexa 647, JF646) lasers. A 63x (NA 1.4, Plan Apochromat DIC) objective was used for imaging. At least 3 positions per condition were selected based on DAPI signal, subsequently automatic multi position imaging was performed and for every position. Fluorescence based autofocus was used to find the center of the nuclei. A z-stack of 11 slices with 500 nm axial spacing from the center was acquired, while the lateral pixel size was 132*132 nm.

### Foci quantification

BRCA2 and RAD51 foci were automatically quantified using CellProfiler (Carpenter, 2006). The analysis script can be found at https://github.com/maartenpaul/DBD_foci. In short, from maximum projections of the confocal images, nuclei were segmented using a global threshold (minimum cross-entropy) based on the DAPI signal. Subsequently within the masked image based on segmented nuclei RAD51 foci were identified using global threshold (Robust background) method with 2 standard deviations above background. The integrated intensity of EdU signal per nucleus was also measured and used to determine the EdU positive cells. Based on the distribution of the integrated intensity of EdU signal per nucleus a fixed threshold was set at 500 a.u., cells above this threshold were defined EdU positive. Also the integrated intensity per focus for BRCA2 and RAD51 were obtained from CellProfiler. The data was exported as CSV files from CellProfiler. R and Rstudio was used to plot the data (example script can be found at the Github repository mentioned above).

### Live cell imaging

For tracking experiments cells were labelled with 5 nM JF549-HaloTag ligand for 15-30 minutes at 37 °C in mouse ES imaging medium (FluoroBrite DMEM (ThermoFisher, 10 % FCS supplemented with nonessential amino acids, 0.1 mM β-mercaptoethanol, pen/strap and 1,000 U/ml leukemia inhibitory factor). Subsequently cells were incubated twice for 15 minutes with fresh imaging medium, while washing the cells once with PBS in between. Microscopy experiments were performed at a Zeiss Elyra PS complemented with a temperature-controlled stage and objective heating (TokaiHit). Samples were kept at 37 °C and 5% CO_2_ while imaging. For excitation of JF549 a 100mW 561 nm laser was used. The samples were illuminated with HiLo illumination by using a 100x 1.57NA Korr αPlan Apochromat (Zeiss) TIRF objective. Andor iXon DU897 was used for detection of the fluorescence signal, from the chip a region of 256 by 256 pixels (with an effective pixel size of 100*100 nm) was recorded at 31.25 Hz interval (30 ms integration time plus 2 ms image transfer time). EMCCD gain was set at 300. Per cell a total of 2000 frames were recorded.

### Single-molecule tracking analysis

Recorded images were converted from LSM-format (Zeiss) to tiff in Fiji (Schindelin et al., 2012) using the Bioformats plugin and prepared for localization and tracking analysis with the SOS plugin (Reuter et al., 2014, http://smal.ws/wp/software/sosplugin/). To track only molecules within the nucleus, for every movie a mask was manually drawn around the nucleus of the cell. A fixed intensity threshold was used to identify molecules in individual frames. The localized molecules were linked through nearest neighbor linking with a maximum displacement of 1.2 μm and a gap size of maximum 1. Tracks had to be at least 5 frames long to be processed further.

Subsequently, the track data was imported in R for analysis using a home-build script (https://github.com/maartenpaul/DBD_tracking). Tracks were segmented in tracklets using the ML-MSS software described in (Arts et al., 2019) (https://github.com/ismal/DL-MSS), using a 3-state deeplearning prediction model. Apparent diffusion constants for the tracklets were estimated by determining the slope of the MSD(t) curve. From all tracklets that were at least 10 frames in length.

### Protein expression and purification

Full-length BRCA2 construct in pHCMV1 was a generous gift from S. Kowalczykowski. Various variants of BRCA2 (BRCA2 ΔDBD, BRCA2 ΔCTD and BRCA2 ΔDBDΔCTD) (**Figure 4A**), with 2 tandem N-terminal maltose binding protein (MBP) tag, were prepared by Q5^®^ site directed mutagenesis (NEB) (**Figure 4A**). Purified plasmids were transfected with 10 % (v/v) PEI transfection solution in 293T HEK cells, adapted for suspension culture, in FreeStyle™ 293 Expression Medium (Gibco) at approximately 10^6^ cells/ml. Transfection solution was prepared by adding 1 μg/ml purified DNA and 2 μg/ml linear PEI in Serum- Free Hybridoma Media (Gibco^®^) supplemented with 1 % FCS. Transfection solution was incubated for 20 minutes at room temperature and added to 500 ml of HEK cell suspension growing at 37 °C with shaking at 250 rpm. After 48 hours, at a cell count of about 2 × 10^6^ /ml, cells were harvested by centrifugation at 8000 × g, 4 °C for 15 minutes. The cell pellet was resuspended in 10 ml ice cold PBS and frozen in liquid nitrogen. Next, cells were lysed in 200 ml lysis buffer (50 mM HEPES pH 7.5, 250 mM NaCl, 1 % NP-40, 1 mM ATP, 3 mM MgCl2, 1 mM Pefabloc SC, 2 tablets EDTA free protease inhibitor (Roche), 1 mM DTT) for 15 minutes at 4 °C with shaking. The lysate was centrifuged at 10,000 × g, 4 °C for 15 minutes. The supernatant was incubated O/N with 10 ml Amylose resin pre-equilibrated in wash buffer (50 mM HEPES pH 7.5, 250 mM NaCl, 0.5 mM EDTA, 1 mM DTT). Next day, the beads were washed 3 times with wash buffer by centrifugation at 2000 × g at 4 °C for 5 minutes and aspiration of the supernatant. The washed resin was incubated with elution buffer (50 mM maltose, 50mM HEPES pH 8.2, 250 mM NaCl, 0.5 mM EDTA, 10 % glycerol, 1 mM DTT, 1 mM Pefabloc SC) for 15 minutes at 4 °C on a rolling platform. The eluate was collected by passing the slurry through a disposable BioRad column at 4 °C. The eluate was loaded on a 1ml HiTrap-Q column from GE using Q low buffer (50 mM HEPES pH 8.2, 250 mM NaCl, 0.5 mM EDTA, 10 % glycerol, 1 mM DTT, 1 mM PMSF) and eluted with Q high buffer (50 mM HEPES pH 8.2, 1 M NaCl, 0.5 mM EDTA, 10 % glycerol, 1 mM DTT, 1 mM PMSF). Peak elution fractions were checked by western blot using anti-BRCA2 antibodies. Fractions with proteins were aliquoted into single use aliquots by snap freezing in liquid nitrogen and stored at −80 °C.

Untagged human RAD51 was expressed and purified as described (Modesti et al., 2007).

### SFM sample preparation, imaging and analyses

For BRCA2-RAD51 reactions, aliquots of BRCA2 stored at −80 °C were thawed and diluted four-fold in 10 mM HEPES pH 8.0 buffer, to subsequently prepare a reaction of 2.5 nM BRCA2 construct in 22 mM HEPES pH 8.2, 112 mM NaCl, 0.125 mM EDTA, 2.5 % glycerol, 0.25 mM DTT. Samples were incubated at 37 °C in the absence or presence of 250 nM RAD51 for 30 minutes without shaking.

For BRCA2-ssDNA reactions, after dilution as mentioned above, the protein was incubated at 37 °C for 30 minutes with linear 90 nt ssDNA oligo (3.4 μM in nt) (5’- AF647/AATTCTCATTTTACTTACCGGACGCTATTAGCAGTGGCAGATTGTACTGAGAGTGCACCATATGCGGTGTGAAATACCGCACAGATGCGT-3’). After incubation 50 μM spermidine was added to the sample.

Samples for SFM imaging were prepared by depositing 20 μl reaction volume on a freshly cleaved mica (Muscovite mica, V5 quality, EMS) for 2 minutes, followed by a 2 ml wash using 18 MΩ water and drying in filtered (0.22 μm) air. SFM images were obtained with a Nanoscope IV (Bruker), using tapping mode in air with a silicon probe, NHC-W, with tip radius <10 nm and resonance frequency range of 310-372 kHz (Nanosensor, Veeco Instruments, Europe). All images were acquired with a scan size of 2 × 2 μm at 512 × 512 pixels per image at 0.5 Hz. Images were processed using Nanoscope analysis (Bruker) for background flattening. Quantitative analysis of the images was performed as described using SFMetric software (Sanchez et al., 2017, Sidhu et al. 2020). In volumetric analyses, a comparison of the oligomeric volume of the different regions with RAD51 (56 nm^3^) showed that the monomer volume of RAD51 is much lower than the threshold volume and thus free RAD51 is removed from analysis (**Figure S2**).

The conformation of the molecules was quantified by the parameters of solidity. Solidity measures the irregular shape of the selected molecule by using the ratio of the area of the selected molecule to the area of a convex hull, that completely encloses the molecule. Solidity is presented in a scale of 1 to 0, where a value of ~1 signifies a globular molecule while a value ~0 represents a highly irregular molecular shape.

## Supporting information

Figure supplements related to main figures

## Acknowledgments

We thank the Optical Imaging Centre for use and technical assistance of the optical microscopes; Ihor Smal (Erasmus MC) for assistance in single-molecule tracking analysis; Luke Lavis (HHMI Janelia) for providing HaloTag ligands; Niklas Bachmann for assistance in making the BRCA2-Halo ΔCTD cell line. We thank Joyce Lebbink (Erasmus MC) and Nick van der Zon (Erasmus MC) for critically reading the manuscript.

## References

Al Abo, M., Dejsuphong, D., Hirota, K., Yonetani, Y., Yamazoe, M., Kurumizaka, H., & Takeda, S. (2014). Compensatory Functions and Interdependency of the DNA-Binding Domain of BRCA2 with the BRCA1-PALB2-BRCA2 Complex. Cancer Research, 74, 797–807. doi:10.1158/0008-5472.CAN-13-1443

Arts, M., Smal, I., Paul, M. W., Wyman, C., & Meijering, E. (2019). Particle Mobility Analysis Using Deep Learning and the Moment Scaling Spectrum. Sci Rep, 9(1), 17160. doi:10.1038/s41598-019-53663-8

Atanassov, B. S., Barrett, J. C., & Davis, B. J. (2005). Homozygous germ line mutation in exon 27 of murine Brca2 disrupts the Fancd2-Brca2 pathway in the homologous recombination-mediated DNA interstrand cross-links’ repair but does not affect meiosis. Genes Chromosomes Cancer, 44(4), 429–437. doi:10.1002/gcc.20255

Baumann, P., & West, S. C. (1998). Role of the human RAD51 protein in homologous recombination and double-stranded-break repair. Trends Biochem Sci, 23(7), 247–251. doi:10.1016/s0968-0004(98)01232-8

Chen, C. F., Chen, P. L., Zhong, Q., Sharp, Z. D., & Lee, W. H. (1999). Expression of BRC repeats in breast cancer cells disrupts the BRCA2-Rad51 complex and leads to radiation hypersensitivity and loss of G(2)/M checkpoint control. J Biol Chem, 274(46), 32931–32935. doi:10.1074/jbc.274.46.32931

Donoho, G., Brenneman, M. A., Cui, T. X., Donoviel, D., Vogel, H., Goodwin, E. H., Chen, D. J., & Hasty, P. (2003). Deletion of Brca2 exon 27 causes hypersensitivity to DNA crosslinks, chromosomal instability, and reduced life span in mice. Genes Chromosomes Cancer, 36(4), 317–331. doi:10.1002/gcc.10148

Dunker, A. K., Cortese, M. S., Romero, P., Iakoucheva, L. M., & Uversky, V. N. (2005). Flexible nets. The roles of intrinsic disorder in protein interaction networks. FEBS Journal, 272(20), 5129–5148. doi:10.1111/j.1742-4658.2005.04948.x

Edwards, S. L., Brough, R., Lord, C. J., Natrajan, R., Vatcheva, R., Levine, D. A., Boyd, J., Reis-Filho, J. S., & Ashworth, A. (2008). Resistance to therapy caused by intragenic deletion in BRCA2. Nature, 451, 1111–1115. doi:10.1038/nature06548

Ehlen, A., Martin, C., Miron, S., Julien, M., Theillet, F. X., Ropars, V., Sessa, G., Beaurepere, R., Boucherit, V., Duchambon, P., El Marjou, A., Zinn-Justin, S., & Carreira, A. (2020). Proper chromosome alignment depends on BRCA2 phosphorylation by PLK1. Nat Commun, 11(1), 1819. doi:10.1038/s41467-020-15689-9

Esashi, F., Christ, N., Cannon, J., Liu, Y., Hunt, T., Jasin, M., & West, S. C. (2005). CDK-dependent phosphorylation of BRCA2 as a regulatory mechanism for recombinational repair. Nature, 434, 598–604. doi:10.1038/nature03404

Essers, J., Hendriks, R. W., Swagemakers, S. M., Troelstra, C., de Wit, J., Bootsma, D., Hoeijmakers, J. H., & Kanaar, R. (1997). Disruption of mouse RAD54 reduces ionizing radiation resistance and homologous recombination. Cell, 89(2), 195–204. doi:10.1016/s0092-8674(00)80199-3

Essers, J., Theil, A. F., Baldeyron, C., van Cappellen, W. A., Houtsmuller, A. B., Kanaar, R., & Vermeulen, W. (2005). Nuclear dynamics of PCNA in DNA replication and repair. Mol Cell Biol, 25(21), 9350–9359. doi:10.1128/MCB.25.21.9350-9359.2005

Feng, W., & Jasin, M. (2017). BRCA2 suppresses replication stress-induced mitotic and G1 abnormalities through homologous recombination. Nature Communications, 8, 525. doi:10.1038/s41467-017-00634-0

Galkin, V. E., Esashi, F., Yu, X., Yang, S., West, S. C., & Egelman, E. H. (2005). BRCA2 BRC motifs bind RAD51-DNA filaments. Proc Natl Acad Sci U S A, 102(24), 8537–8542. doi:10.1073/pnas.0407266102

Ghouil, R., Miron, S., Koornneef, L., Veerman, J., Paul, M. W., Du, M.-H. L., Sleddens-Linkels, E., Rossum-Fikkert, S. E. V., Loon, Y. v., Felipe-Medina, N., Pendas, A. M., Maas, A., Essers, J., Legrand, P., Baarends, W. M., Kanaar, R., Zinn-Justin, S., & Zelensky, A. N. (2020). A cryptic BRCA2 repeated motif binds to HSF2BP oligomers with no impact on meiotic recombination. bioRxiv, 2020.2012.2029.424679. doi:10.1101/2020.12.29.424679

Gudmundsdottir, K., Lord, C. J., Witt, E., Tutt, A. N., & Ashworth, A. (2004). DSS1 is required for RAD51 focus formation and genomic stability in mammalian cells. EMBO Rep, 5(10), 989–993. doi:10.1038/sj.embor.7400255

Gunasekaran, K., Tsai, C. J., Kumar, S., Zanuy, D., & Nussinov, R. (2003). Extended disordered proteins: targeting function with less scaffold. Trends Biochem Sci, 28(2), 81–85. doi:10.1016/S0968-0004(03)00003-3

Hartford, S. A., Chittela, R., Ding, X., Vyas, A., Martin, B., Burkett, S., Haines, D. C., Southon, E., Tessarollo, L., & Sharan, S. K. (2016). Interaction with PALB2 Is Essential for Maintenance of Genomic Integrity by BRCA2. PLoS Genet, 12(8), e1006236. doi:10.1371/journal.pgen.1006236

Heyer, W. D., Ehmsen, K. T., & Liu, J. (2010). Regulation of homologous recombination in eukaryotes. Annu Rev Genet, 44, 113–139. doi:10.1146/annurev-genet-051710-150955

Holloman, W. K. (2011). Unraveling the mechanism of BRCA2 in homologous recombination. Nat Struct Mol Biol, 18(7), 748–754. doi:10.1038/nsmb.2096

Hooper, M., Hardy, K., Handyside, A., Hunter, S., & Monk, M. (1987). HPRT-deficient (Lesch-Nyhan) mouse embryos derived from germline colonization by cultured cells. Nature, 326(6110), 292–295. doi:10.1038/326292a0

Jensen, R. B., Carreira, A., & Kowalczykowski, S. C. (2010). Purified human BRCA2 stimulates RAD51-mediated recombination. Nature, 467(7316), 678–683. doi:nature09399 [pii] 10.1038/nature09399

Kojic, M., Zhou, Q., Lisby, M., & Holloman, W. K. (2005). Brh2-Dss1 interplay enables properly controlled recombination in Ustilago maydis. Mol Cell Biol, 25(7), 2547–2557. doi:10.1128/MCB.25.7.2547-2557.2005

Le, H. P., Ma, X., Vaquero, J., Brinkmeyer, M., Guo, F., Heyer, W. D., & Liu, J. (2020). DSS1 and ssDNA regulate oligomerization of BRCA2. Nucleic Acids Res. doi:10.1093/nar/gkaa555

Li, J., Zou, C., Bai, Y., Wazer, D. E., Band, V., & Gao, Q. (2006). DSS1 is required for the stability of BRCA2. Oncogene, 25(8), 1186–1194. doi:10.1038/sj.onc.1209153

Lindenburg, L. H., Pantelejevs, T., Gielen, F., Zuazua-Villar, P., Butz, M., Rees, E., Kaminski, C. F., Downs, J. A., Hyvönen, M., & Hollfelder, F. (2020). Improved RAD51 binders through motif shuffling based on the modularity of BRC repeats. bioRxiv.

Liu, J., Doty, T., Gibson, B., & Heyer, W. D. (2010). Human BRCA2 protein promotes RAD51 filament formation on RPA-covered single-stranded DNA. Nat Struct Mol Biol, 17(10), 1260–1262. doi:nsmb.1904 [pii] 10.1038/nsmb.1904

Lomonosov, M., Anand, S., Sangrithi, M., Davies, R., & Venkitaraman, A. R. (2003). Stabilization of stalled DNA replication forks by the BRCA2 breast cancer susceptibility protein. Genes Dev, 17(24), 3017–3022. doi:10.1101/gad.279003

Los, G. V., Encell, L. P., McDougall, M. G., Hartzell, D. D., Karassina, N., Zimprich, C., Wood, M. G., Learish, R., Ohana, R. F., Urh, M., Simpson, D., Mendez, J., Zimmerman, K., Otto, P., Vidugiris, G., Zhu, J., Darzins, A., Klaubert, D. H., Bulleit, R. F., & Wood, K. V. (2008). HaloTag: a novel protein labeling technology for cell imaging and protein analysis. ACS Chem Biol, 3(6), 373–382. doi:10.1021/cb800025k

Marple, T., Kim, T. M., & Hasty, P. (2006). Embryonic stem cells deficient for Brca2 or Blm exhibit divergent genotoxic profiles that support opposing activities during homologous recombination. Mutat Res, 602(1-2), 110–120. doi:10.1016/j.mrfmmm.2006.08.005

Morimatsu, M., Donoho, G., & Hasty, P. (1998). Cells deleted for Brca2 COOH terminus exhibit hypersensitivity to gamma-radiation and premature senescence. Cancer Res, 58(15), 3441–3447.

Naipal, K. A., Verkaik, N. S., Ameziane, N., van Deurzen, C. H., Ter Brugge, P., Meijers, M., Sieuwerts, A. M., Martens, J. W., O’Connor, M. J., Vrieling, H., Hoeijmakers, J. H., Jonkers, J., Kanaar, R., de Winter, J. P., Vreeswijk, M. P., Jager, A., & van Gent, D. C. (2014). Functional ex vivo assay to select homologous recombination-deficient breast tumors for PARP inhibitor treatment. Clin Cancer Res, 20(18), 4816–4826. doi:10.1158/1078-0432.CCR-14-0571

Oliver, A. W., Swift, S., Lord, C. J., Ashworth, A., & Pearl, L. H. (2009). Structural basis for recruitment of BRCA2 by PALB2. EMBO Rep, 10(9), 990–996. doi:embor2009126 [pii] 10.1038/embor.2009.126

Pellegrini, L., Yu, D. S., Lo, T., Anand, S., Lee, M., Blundell, T. L., & Venkitaraman, A. R. (2002). Insights into DNA recombination from the structure of a RAD51-BRCA2 complex. Nature, 420(6913), 287–293. doi:10.1038/nature01230 nature01230 [pii]

Prakash, R., Zhang, Y., Feng, W., & Jasin, M. (2015). Homologous recombination and human health: the roles of BRCA1, BRCA2, and associated proteins. Cold Spring Harb Perspect Biol, 7(4), a016600. doi:10.1101/cshperspect.a016600

Ran, F. A., Hsu, P. D., Wright, J., Agarwala, V., Scott, D. A., & Zhang, F. (2013). Genome engineering using the CRISPR-Cas9 system. Nat Protoc, 8(11), 2281–2308. doi:10.1038/nprot.2013.143

Reuter, M., Zelensky, A., Smal, I., Meijering, E., van Cappellen, W. A., de Gruiter, H. M., van Belle, G. J., van Royen, M. E., Houtsmuller, A. B., Essers, J., Kanaar, R., & Wyman, C. (2014). BRCA2 diffuses as oligomeric clusters with RAD51 and changes mobility after DNA damage in live cells. The Journal of Cell Biology, 207, 599–613. doi:10.1083/jcb.201405014

Sanchez, H., Paul, M. W., Grosbart, M., van Rossum-Fikkert, S. E., Lebbink, J. H. G., Kanaar, R., Houtsmuller, A. B., & Wyman, C. (2017). Architectural plasticity of human BRCA2-RAD51 complexes in DNA break repair. Nucleic Acids Res, 45(8), 4507–4518. doi:2972667 [pii] 10.1093/nar/gkx084

Sarkisian, C. J., Master, S. R., Huber, L. J., Ha, S. I., & Chodosh, L. A. (2001). Analysis of murine Brca2 reveals conservation of protein-protein interactions but differences in nuclear localization signals. J Biol Chem, 276(40), 37640–37648. doi:10.1074/jbc.M106281200

Schlacher, K., Christ, N., Siaud, N., Egashira, A., Wu, H., & Jasin, M. (2011). Double-strand break repair-independent role for BRCA2 in blocking stalled replication fork degradation by MRE11. Cell, 145, 529–542. doi:10.1016/j.cell.2011.03.041

Shahid, T., Soroka, J., Kong, E., Malivert, L., McIlwraith, M. J., Pape, T., West, S. C., & Zhang, X. (2014). Structure and mechanism of action of the BRCA2 breast cancer tumor suppressor. Nat Struct Mol Biol, 21(11), 962–968. doi:10.1038/nsmb.2899

Sharan, S. K., Morimatsu, M., Albrecht, U., Lim, D.-S., Regel, E., Dinh, C., Sands, A., Eichele, G., Hasty, P., & Bradley, A. (1997). Embryonic lethality and radiation hypersensitivity mediated by Rad51 in mice lacking Brca2. Nature, 386, 804–810. doi:10.1038/386804a0

Shcherbakova, D. M., & Verkhusha, V. V. (2013). Near-infrared fluorescent proteins for multicolor in vivo imaging. Nat Methods, 10(8), 751–754. doi:10.1038/nmeth.2521

Siaud, N., Barbera, M. A., Egashira, A., Lam, I., Christ, N., Schlacher, K., Xia, B., & Jasin, M. (2011). Plasticity of BRCA2 function in homologous recombination: Genetic interactions of the PALB2 and DNA binding domains. PLoS Genetics, 7. doi:10.1371/journal.pgen.1002409

Sidhu, A., Grosbart, M., Sanchez, H., Verhagen, B., van der Zon, N. L. L., Ristic, D., van Rossum-Fikkert, S. E., & Wyman, C. (2020). Conformational flexibility and oligomerization of BRCA2 regions induced by RAD51 interaction. Nucleic Acids Res. doi:10.1093/nar/gkaa648

Spain, B. H., Larson, C. J., Shihabuddin, L. S., Gage, F. H., & Verma, I. M. (1999). Truncated BRCA2 is cytoplasmic: implications for cancer-linked mutations. Proc Natl Acad Sci U S A, 96(24), 13920–13925. doi:10.1073/pnas.96.24.13920

van der Lee, R., Buljan, M., Lang, B., Weatheritt, R. J., Daughdrill, G. W., Dunker, A. K., Fuxreiter, M., Gough, J., Gsponer, J., Jones, D. T., Kim, P. M., Kriwacki, R. W., Oldfield, C. J., Pappu, R. V., Tompa, P., Uversky, V. N., Wright, P. E., & Babu, M. M. (2014). Classification of intrinsically disordered regions and proteins. Chem Rev, 114(13), 6589–6631. doi:10.1021/cr400525m

Whelan, D. R., Lee, W. T. C., Yin, Y., Ofri, D. M., Bermudez-Hernandez, K., Keegan, S., Fenyo, D., & Rothenberg, E. (2018). Spatiotemporal dynamics of homologous recombination repair at single collapsed replication forks. Nat Commun, 9(1), 3882. doi:10.1038/s41467-018-06435-3

Xia, B., Dorsman, J. C., Ameziane, N., de Vries, Y., Rooimans, M. A., Sheng, Q., Pals, G., Errami, A., Gluckman, E., Llera, J., Wang, W., Livingston, D. M., Joenje, H., & de Winter, J. P. (2007). Fanconi anemia is associated with a defect in the BRCA2 partner PALB2. Nat Genet, 39(2), 159–161. doi:10.1038/ng1942

Xia, B., Sheng, Q., Nakanishi, K., Ohashi, A., Wu, J., Christ, N., Liu, X., Jasin, M., Couch, F. J., & Livingston, D. M. (2006). Control of BRCA2 cellular and clinical functions by a nuclear partner, PALB2. Mol Cell, 22(6), 719–729. doi:10.1016/j.molcel.2006.05.022

Yang, H., Jeffrey, P. D., Miller, J., Kinnucan, E., Sun, Y., Thoma, N. H., Zheng, N., Chen, P. L., Lee, W. H., & Pavletich, N. P. (2002). BRCA2 function in DNA binding and recombination from a BRCA2-DSS1-ssDNA structure. Science, 297(5588), 1837–1848. doi:10.1126/science.297.5588.1837297/5588/1837 [pii]

Yao, X., Wang, X., Hu, X., Liu, Z., Liu, J., Zhou, H., Shen, X., Wei, Y., Huang, Z., Ying, W., Wang, Y., Nie, Y. H., Zhang, C. C., Li, S., Cheng, L., Wang, Q., Wu, Y., Huang, P., Sun, Q., Shi, L., & Yang, H. (2017). Homology-mediated end joining-based targeted integration using CRISPR/Cas9. Cell Res, 27(6), 801–814. doi:10.1038/cr.2017.76

Yu, V. P., Koehler, M., Steinlein, C., Schmid, M., Hanakahi, L. A., van Gool, A. J., West, S. C., & Venkitaraman, A. R. (2000). Gross chromosomal rearrangements and genetic exchange between nonhomologous chromosomes following BRCA2 inactivation. Genes Dev, 14(11), 1400–1406.

Yuan, S. S., Lee, S. Y., Chen, G., Song, M., Tomlinson, G. E., & Lee, E. Y. (1999). BRCA2 is required for ionizing radiation-induced assembly of Rad51 complex in vivo. Cancer Res, 59(15), 3547–3551.

Zelensky, A. N., Schimmel, J., Kool, H., Kanaar, R., & Tijsterman, M. (2017). Inactivation of Pol theta and C-NHEJ eliminates off-target integration of exogenous DNA. Nat Commun, 8(1), 66. doi:10.1038/s41467-017-00124-3

Zhao, W., Vaithiyalingam, S., San Filippo, J., Maranon, D. G., Jimenez-Sainz, J., Fontenay, G. V., Kwon, Y., Leung, S. G., Lu, L., Jensen, R. B., Chazin, W. J., Wiese, C., & Sung, P. (2015). Promotion of BRCA2-Dependent Homologous Recombination by DSS1 via RPA Targeting and DNA Mimicry. Mol Cell, 59(2), 176–187. doi:10.1016/j.molcel.2015.05.032

