## Supplementary material for "Role of BRCA2 DNA-binding and C-terminal domain on its mobility and conformation in DNA repair": Figure supplements related to main figures

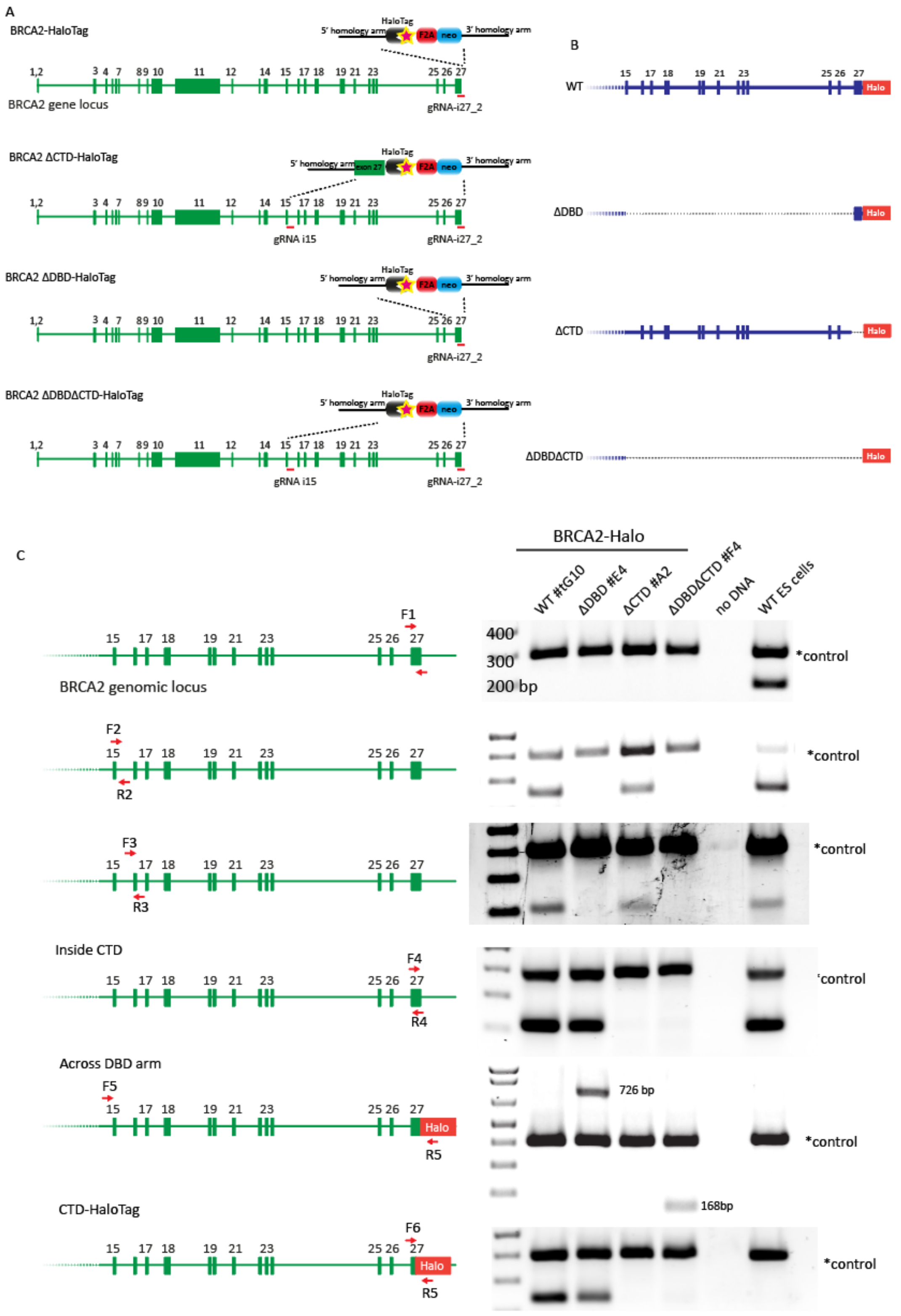


**Figure 1 – supplement 1. Generation of different variant of BRCA2-HaloTag knock-ins** (A) Cartoon depicting the CRISPR/Cas9 targetting strategy, indicating the approximate position of the gRNA sequences (**Supplemental table S1**) and the targetting construct with homology arm and targetted cassette. (B) Scheme displaying the expected BRCA2 locus after targeting the different BRCA2 deletion variants, showing which exons and introns are deleted. (C) Genotyping PCRs to validate the different cell lines. The multiplex PCR reaction also included primers annealing outside the targetted genomic region to serve as a positive control for genomic DNA amplification efficiency. Primers sequences are reported in **Supplemental table S2**.

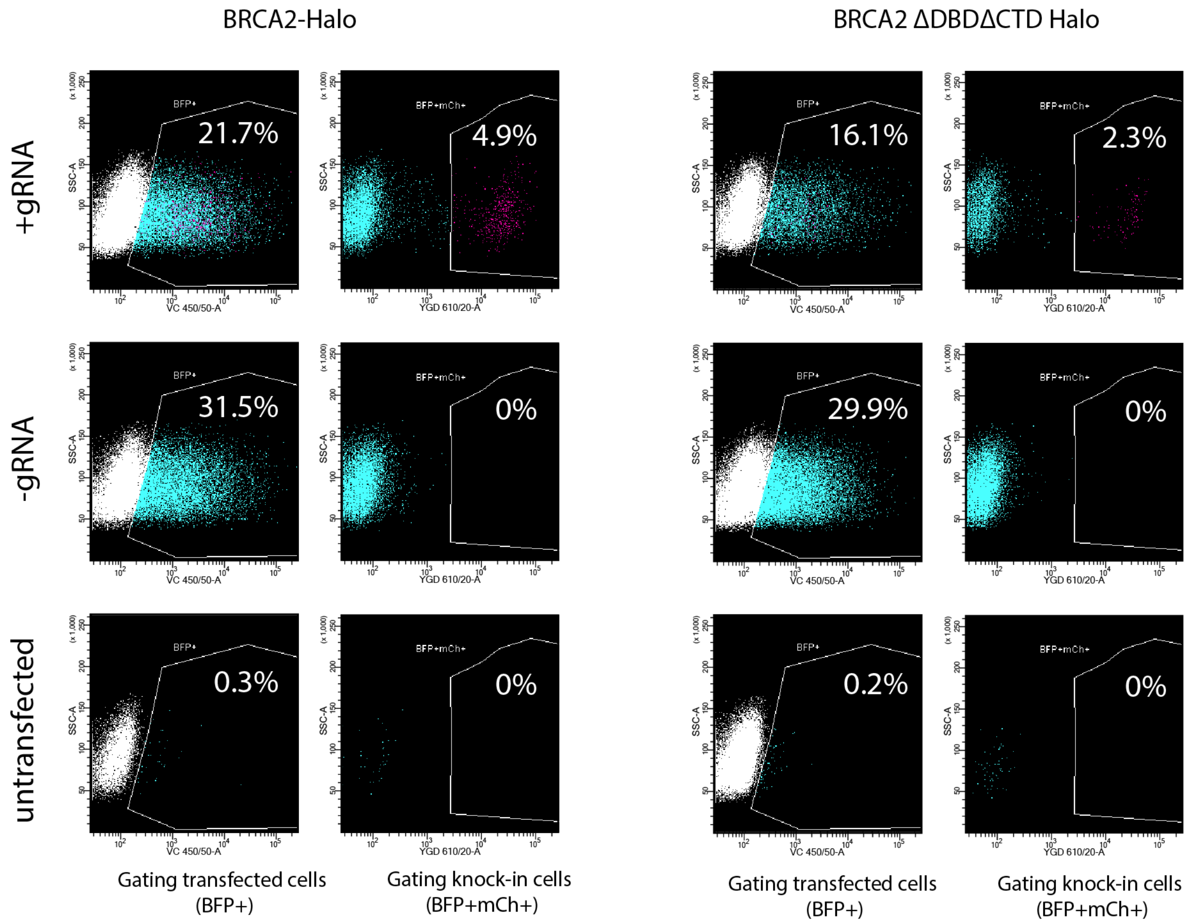


**Figure 1 – supplement 2.** Flow cytometry analysis of the CRISPR/Cas9 βActin-P2A-mCherry targeting assay as shown in **Figure 1G**. A plasmid expressing BFP2 was co-introduced for the selection of successfully transfected cells. The percentage of transfected cells (BFP+) that express mCherry transfected cells was quantified. Typical results shown for full-length BRCA2-Halo and for BRCA2-Halo ΔDBDΔCTD.

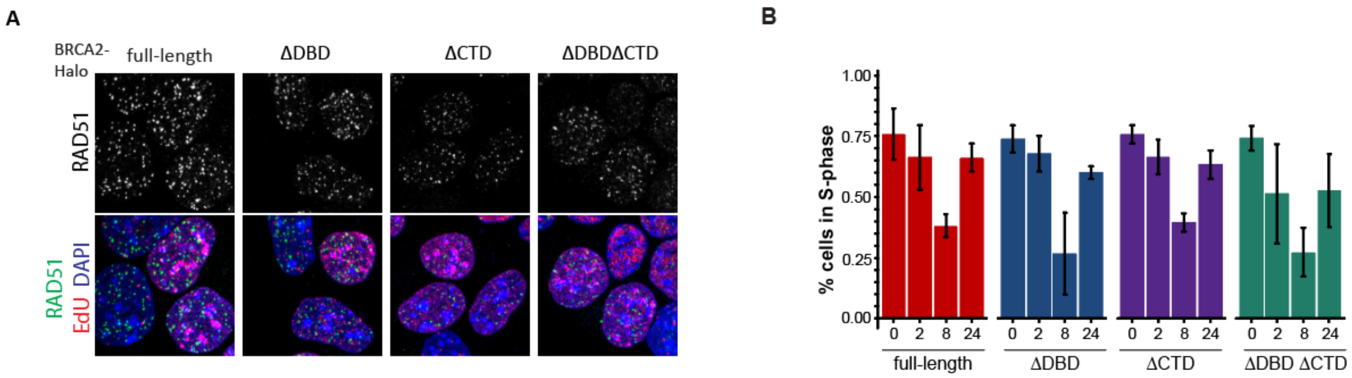


**Figure 2 - Supplement 1** (A) Confocal images (maximum intensity projection) of mES cells producing the indicated variants of BRCA2 immunostained for RAD51 after pre-extraction and pulse-labeling with EdU click chemistry to reveal S-phase cells. (B) Fraction of EdU-positive (S-phase) mES cells revealed as shown in panel (A) at different time points after irradiation with 2 Gy. Average from 3 experiments, bars indicated S.E.M..

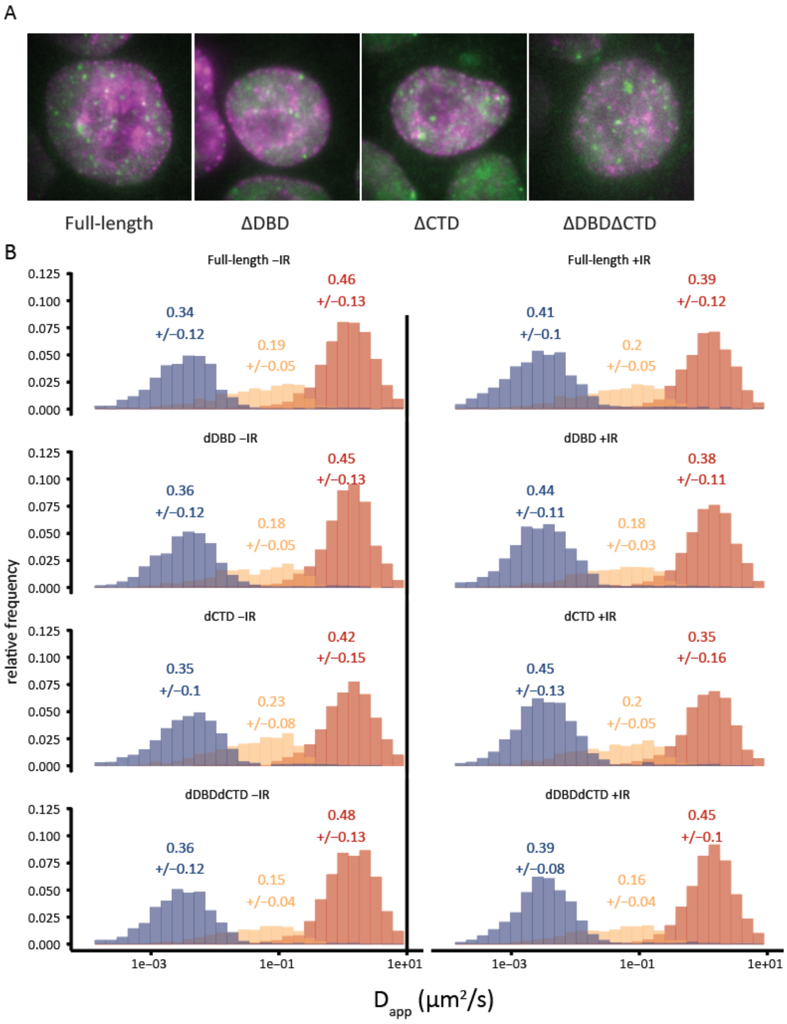


**Figure 3 – Supplement 1** (A) Wide field images of cells from the different BRCA2 variants (like in **Figure 3A**), showing iRFP720-PCNA in magenta and BRCA2-Halo::JF549 in green. (B) Histograms of the apparent diffusion constant estimated for every tracklet. Tracks are segmented in fast (red), slow (yellow) and immobile (blue) tracklets using the DL-MSS software. The numbers above the plots indicate the average fractions +/- standard deviation estimated from fractions per cell.

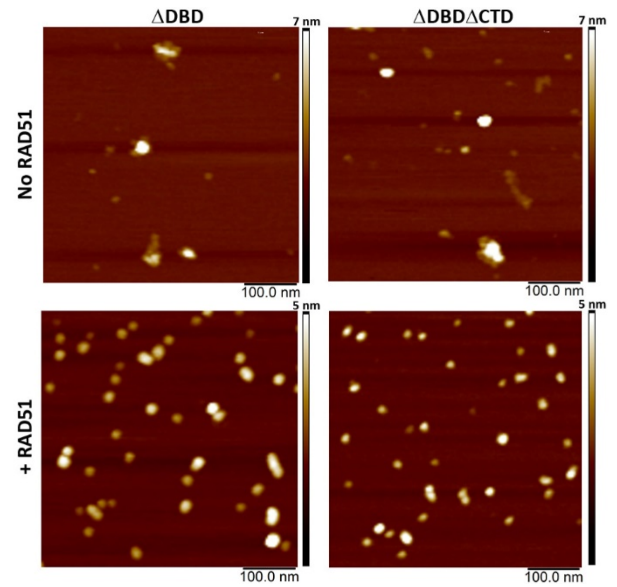


**Figure 4 - supplement 1.** Representative SFM height images of BRCA2 ΔDBD and BRCA2 ΔDBDΔCTD in the presence and absence of RAD51. Both the constructs show reorganization of protein into smaller globular assemblies on interaction with RAD51.

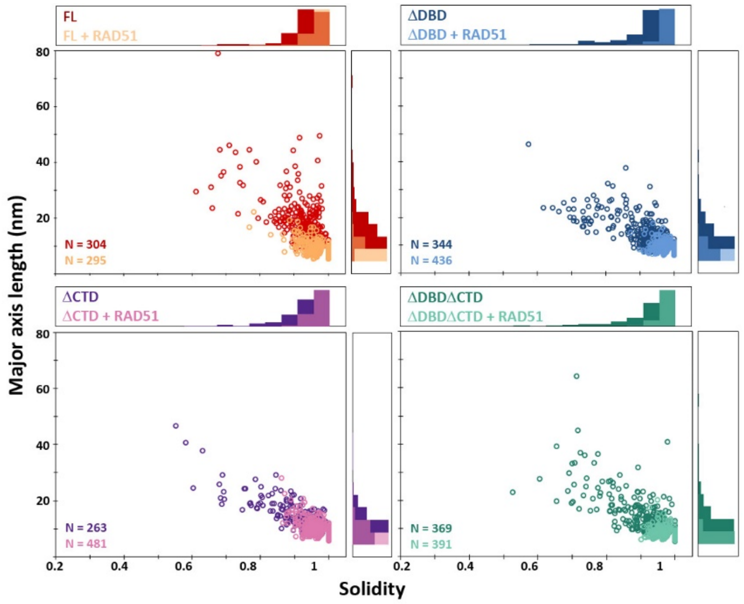


**Figure 4 - supplement 2.** Plots showing distribution of full-lengthBRCA2 and c-terminal deletion constructs in the presence and absence of RAD51. All the BRCA2 samples, full-length and the deletion constructs, show a similar rearrangement into globular and monomeric assemblies.

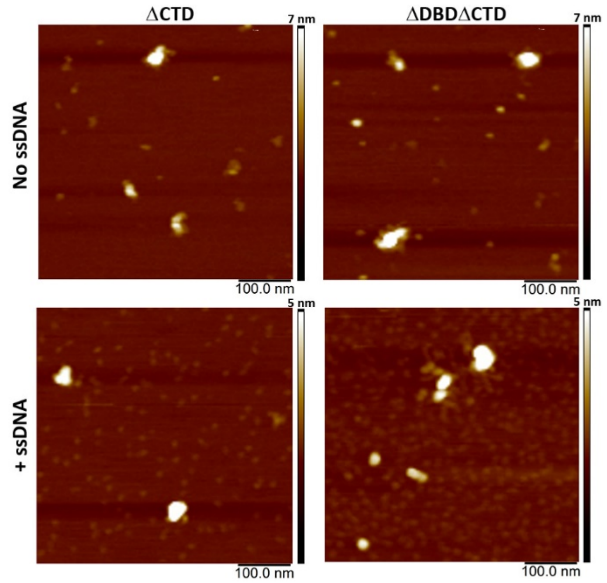


**Figure 5 - supplement 1.** Representative SFM height images of BRCA2 ΔCTD and BRCA2 ΔDBDΔCTD in the presence and absence of ssDNA. Neither of the constructs exhibit a change in conformation on incubation with ssDNA. Noticeable amount of free ssDNA is visible in the background showing lack of interaction.

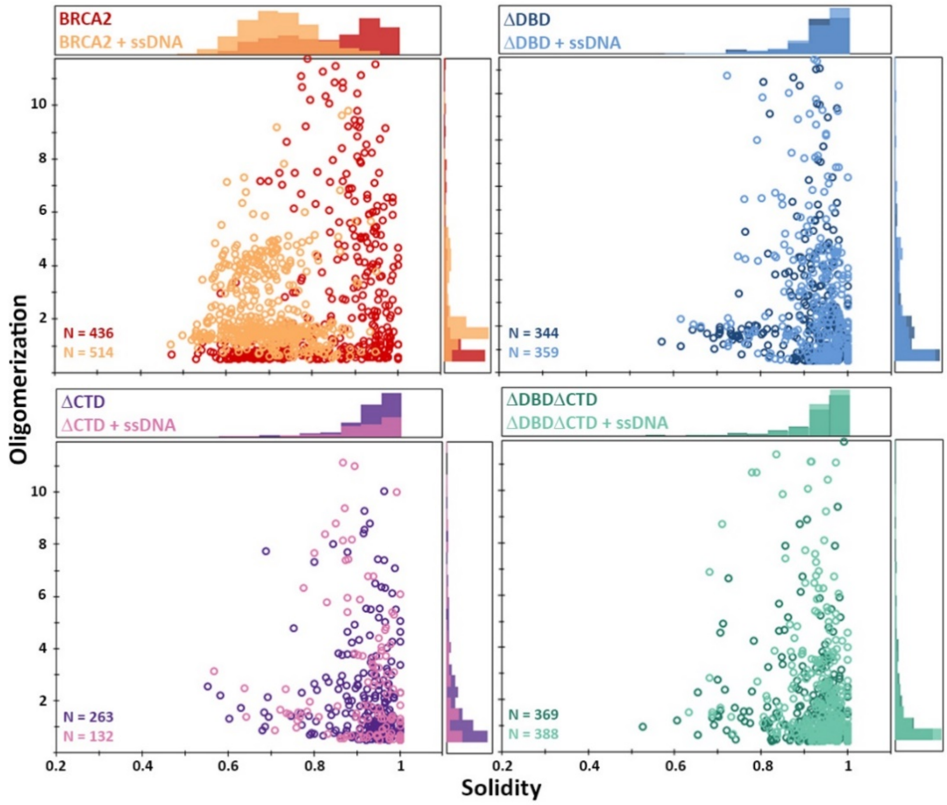


Figure 5 – supplement 2. Distribution of full-length BRCA2 and the C-terminal deletion constructs with respect to their oligomerization and solidity. Distribution of oligomerization and solidity of full-length-BRCA2 and DBD and/or CTD deletion variants. Full-length-BRCA2 rearranges to form extended monomer-dimers and tetramers on interaction with ssDNA, whereas the deletion constructs do not show change in their distribution.
