## Supplemental Figures S1-S2 for "Role of BRCA2 DNA-binding and C-terminal domain on its mobility and conformation in DNA repair"

### SUPPLEMENTARY FIGURES S1-S2

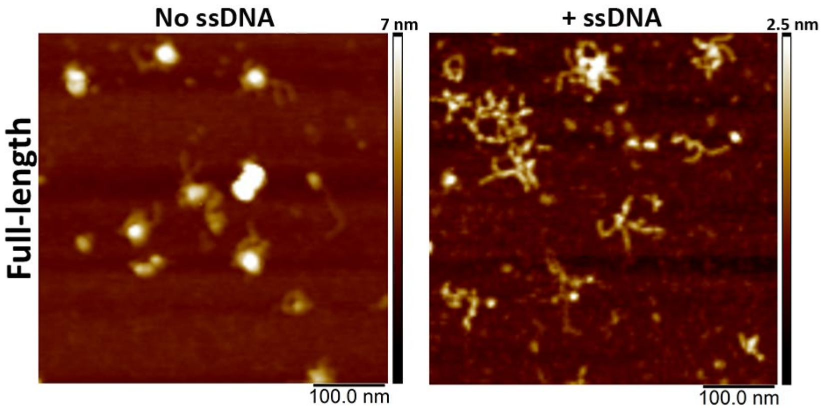


**Figure S1.** Control experiment showing representative SFM height images of full-length BRCA2±ssDNA in the absence of spermidine, showing that the conformational change observed on interaction with ssDNA is not an artifact due to presence of spermidine, which is used to facilitate adsorption of DNA on the mica surface for SFM imaging.

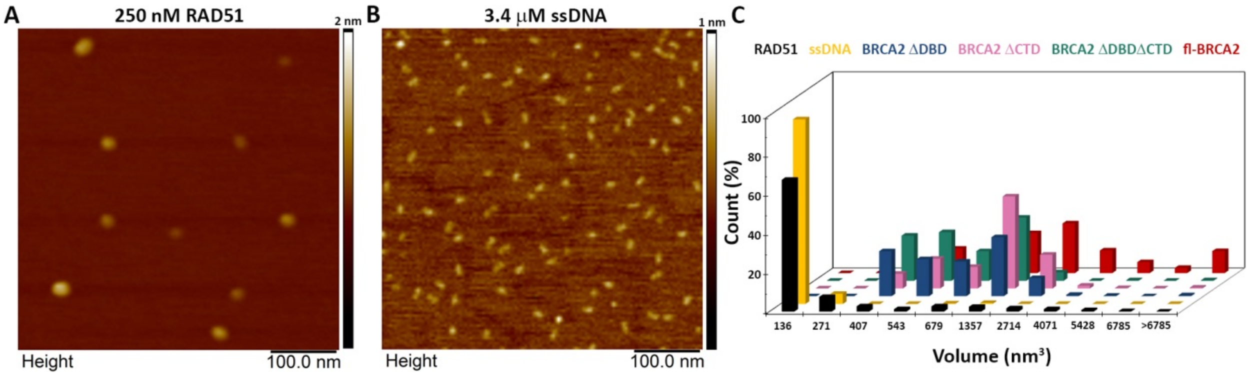


**Figure S2. (A)** Representative SFM height images of RAD51 alone. (B) Representative SFM height image of ssDNA alone. (C) Distribution of volume of all the proteins and ssDNA used in the study.
