## Supplemental Tables S1-S4 for "Role of BRCA2 DNA-binding and C-terminal domain on its mobility and conformation in DNA repair"

### SUPPLEMENTARY TABLES S1-S4

**Table S1. gRNA and their sequences used for generating the BRCA2-Halo knock-in cell lines**

| | gRNA 1 | gRNA 2 |
|---|---|---|
| BRCA2-Halo | gRNA i27_2 | |
| BRCA2-Halo dDBD | gRNA i15 | gRNA i27_2 |
| BRCA2-Halo dCTD | gRNA e27 | gRNA i27_2 |
| BRCA2-Halo dDBDdCTD | gRNA i15 | gRNA i27_2 |

| | Sequence (+PAM) | Target site |
|---|---|---|
| gRNA i27_2 | gctgttgagtcttagcctcc-cgg | across stop-codon |
| gRNA i15 | tgaggcttcccttaggattg-cgg | intron 15 |
| gRNA e27 | ttatcacactgcacactgaa-agg | exon 27 |

**Table S2. Primers used for genotyping the cell lines**

| Name | Alternative name | sequence |
|---|---|---|
| F1 | MP_BRCA2_GT3_R | AACACCTGGCACATAGGTCA |
| R1 | MP_BRCA2_GT4_F | CAGGTGCACAGCAGAGAAGA |
| F2 | MP_DBD_FW | GCAGGCTGCAGTAGGAGACA |
| R2 | MP_BR2_GT6_R | CAGAGCACTTGCCATCAAGA |
| F3 | MP_BR2_GT7_F | ATCACATTTTTCTATTGTTTCTTTTG |
| R3 | MP_BR2_GT7_R | TTCTTCTTTTCCAGCCTTGC |
| F4 | MP_BR2_GT8_F | GGTCAGCAAAGTTATCAAAGTCC |
| R4 | MP_BR2_GT8_R | CTGTGCAGCTGGAGAGACAA |
| F5 | MP_BRCA2_DBD_FIX_FW | GCTGGCCTGGAACTTACTGA |
| R5 | MP_SCRSEQ_HALO_N_R | CACATAATGGGGGTCGAATG |
| F6 | BRCA2-CG-SCRF | TGAAGAACTTGCCTTGCTCA |
| control F | MP_N_BR2_GT1_F | CAGGCTTGTGCAGTGTCTGT |
| control R | MP_N_BR2_GT1_R | TTTTCTGCCTCAGCTTCCAT |

**Table S3. Summary of average solidity of BRCA2 variants in the absence and presence of ssDNA (90-mer) oligomer.**

| BRCA2 variant | 37 °C | + ssDNA |
|---|---|---|
| **Full-length** | 0.82 | 0.69 |
| **ΔDBD** | 0.92 | 0.93 |
| **ΔCTD** | 0.90 | 0.92 |
| **ΔDBDΔCTD** | 0.92 | 0.93 |

**Table S4. Percent rod-like assemblies in BRCA2-RAD51 interaction in various constructs.**

| Complex | Percent rod-like assemblies |
|---|---|
| **Fl-BRCA2-RAD51** | 33 |
| **BRCA2 ΔDBD-RAD51** | 7 |
| **BRCA2 ΔCTD-RAD51** | 40 |
| **BRCA2 ΔDBDΔCTD-RAD51** | 8 |
